# Manufacturing CD20/CD19-targeted iCasp9 regulatable CAR-T_SCM_ cells using *qCART*, the *Quantum pBac*-based CAR-T system

**DOI:** 10.1101/2022.05.03.490475

**Authors:** Peter S. Chang, Yi-Chun Chen, Wei-Kai Hua, Jeff C. Hsu, Jui-Cheng Tsai, Yi-Wun Huang, Yi-Hsin Kao, Pei-Hua Wu, Yi-Fang Chang, Ming-Chih Chang, Yu-Cheng Chang, Shiou-Ling Jian, Jiann-Shiun Lai, Ming-Tain Lai, Wei-Cheng Yang, Chia-Ning Shen, Kuo-Lan Karen Wen, Sareina Chiung-Yuan Wu

**Affiliations:** GenomeFrontier Therapeutics, Inc. Taipei City, Taiwan (R.O.C.); Division of Hematology and Oncology, Department of Internal Medicine, Mackay Memorial Hospital, Taipei, Taiwan (R.O.C.); Department of Medical Research, Laboratory of Good Clinical Research Center, Mackay Memorial Hospital, Tamsui District, New Taipei City, Taiwan (R.O.C.); Department of Medicine, Mackay Medical College, New Taipei City, Taiwan (R.O.C.); OBI Pharma, Inc.; Biomedical Translation Research Center, Academia Sinica, Taipei, Taiwan (R.O.C.); Genomics Research Center, Academia Sinica, Taipei, Taiwan (R.O.C.)

**Keywords:** multiplex CAR-T, *piggyBac*, T_SCM_, transposon

## Abstract

**Background:** CD19-targeted chimeric antigen receptor therapies (CAR19) have driven a paradigm shift in the treatment of relapsed/refractory B-cell malignancies. However, >50% of CAR19-treated patients experienced progressive disease mainly due to antigen escape and low persistence. Clinical prognosis is heavily influenced by CAR-T cell function and systemic cytokine toxicities. Furthermore, it remains a challenge to efficiently, cost-effectively, and consistently manufacture clinically relevant number of virally engineered CAR-T cells.

**Methods:** Using a highly efficient *piggyBac* transposon-based vector, *Quantum pBac*, we developed a virus-free cell engineering system, *Quantum CART (qCART*™), for development and production of multiplex CAR-T therapies.

**Results:** Here, we demonstrated *in vitro and in vivo* that consistent, robust, and functional CD20/CD19 dual-targeted CAR-T stem cell memory (T_SCM_) cells can be efficiently manufactured using the *qCART*™ system for clinical application. *qCART*™-manufactured CAR-T cells from cancer patients expanded efficiently, rapidly eradicated tumors, and can be safely controlled via an iCasp9 suicide gene-inducing drug.

**Conclusions:** The *qCART*™ system is an elegant system for the manufacturing of CAR-T products having all the desired CAR-T therapy attributes. We believe that the simplicity of manufacturing multiplex CAR-T cells using the *qCART*™ system will not only significantly enhance the accessibility of CAR-T therapy but also unlock the full potential of armored CAR-T therapy for the treatment of solid tumors in the future.

**What is already known on this topic:** Despite the considerable success which has been achieved with CD19-targeted chimeric antigen receptor therapies (CAR19), >50% of CAR19-treated patients still experienced progressive disease. Therefore, there is a need to further improve CAR19 therapies. Current CAR19 therapies commonly utilize virus-based cell engineering methods. CAR-T production using these methods face multiple hurdles, including difficulties to efficiently, cost-effectively, and consistently manufacture clinically relevant number of CAR-T cells. We have previously used a highly efficient *piggyBac* transposon-based vector, *Quantum pBac*, to establish *Quantum CART* (*qCART*™) which is a virus-free cell engineering system for development and production of multiplex CAR-T therapies.

**What this study adds:** In this report, we further demonstrate *in vitro* and *in vivo* that consistent, robust, and functional iCasp9-regulatable, CD20/CD19 dual-targeted CAR-T stem cell memory (T_SCM_) cells can be efficiently manufactured using the *qCART*™ system for clinical application. These cells possess all the desired attributes for ensuring therapeutic efficacy in CAR-T therapy, including high CAR-T_SCM_, balanced CD8/CD4 ratio, low exhaustion and senescence marker expressions, and high *ex vivo* and *in vivo* expansion capacity. Importantly, we show that *qCART*™-manufactured CAR-T cells from hematological cancer patients expanded efficiently, effectively eradicated tumors, and can be safely controlled via an iCasp9 suicide gene-inducing drug. We believe that the simplicity of manufacturing multiplex CAR-T cells using the *qCART*™ system will not only significantly enhance the accessibility of CAR-T therapy but also unlock the full potential of armored CAR-T therapy for the treatment of solid tumors in the future.

**How this study might affect research, practice or policy:** Our findings demonstrate that *qCART*™ is a virus-free CAR-T engineering system for manufacturing CAR-T_SCM_ cells from either healthy donors or hematological cancer patients, that possess all the desired attributes for a successful CAR-T therapy. These cells expanded efficiently, rapidly eradicated tumors, and can be safely controlled via activation of iCasp9. We expect that this simple yet robust system for manufacturing multiplex CAR-T cells will advance the CAR-T field.

## BACKGROUND

Over the last decade, chimeric antigen receptor (CAR)-T therapy has become one of the most promising cancer treatment, but the heavy reliance on lentiviral and retroviral vectors for CAR-T cell manufacturing makes the therapy expensive and inaccessible to the masses.[1] Frequent cytokine release syndrome (CRS) and immune effector cell-associated neurotoxicity syndrome (ICANS) associated with clinical patients treated with conventional CAR-T therapies have raised safety concerns.[2] Furthermore, virus-based gene therapies have limited gene payload capacity, making them less suitable for engineering multiplex armored CAR-T cells.[3,4]

To date, B-cell malignancies remain the most actively-studied cancers for CAR-T therapy development. Since *Brentjens et al. (2003)* first demonstrated the eradication of B-cell tumors by CD19-targeted second-generation CAR-T cells, several commercial CAR-T products have been approved by FDA.[5,6] However, treatment with CAR19 therapy resulted in only 30-40% long-term progression-free survival (PFS) in aggressive Non-Hodgkin lymphoma (NHL) patients, and the median event-free survival (EFS) of adult B-cell ALL patients was 6.1 months.[7,8] Evidence suggest that downregulation of CD19 may in part be responsible for patient relapse.[9,10] As demonstrated by recent clinical studies, bispecific CD19/CD20-targeted CAR-T therapy improved patient survival compared to CAR19 therapy.[11,12]

Transposon system, such as *Sleeping Beauty* and *piggyBac*, is a virus-free alternative for generating CAR-T cells but suffers from low gene transfer efficiency and limited expansion of engineered cells.[3,13,14] We recently developed the most advanced version of *piggyBac* known as *Quantum pBac (qPB)* that is 15 times more active than *hyperactive piggyBac* in human T cells.[15,16] We previously developed a novel genetic engineering system known as *Quantum CART (qCART*™) that combines efficient transgene design, evaluation, and identification of lead construct (*GTailor*); virus-free vector (*Quantum pBac*); effective gene delivery (*Quantum Nufect*); and optimized cell expansion (*iCellar*).[15,16] We reported that *qCART*™ could be used to engineer highly potent T stem cell memory (T_SCM_) CAR-Tcells. Here, we further validate that the *qCART*™ system can be used to rapidly manufacture highly potent patient-derived bicistronic CD20/CD19-targeted iCasp9 regulatable CAR-T cells (GF-CART01) for treating B-cell malignancies. We demonstrate enhanced anti-tumor efficacy of GF-CART01 cells both *in vitro* and in a xenotransplant immunodeficient mouse model.

## METHODS

### Cell engineering

CD19-expressing K562 cells were generated by using a *piggyBac*-based cell engineering system. K562 cells were nucleofected using *Quantum Nufect*™ (GenomeFrontier Therapeutics) with a plasmid carrying CD19 and hygromycin genes followed by hygromycin selection for 14 days. To select for clones that homogeneously express CD19, limiting-dilution was carried out to reach a concentration of one cell/well. Following clonal expansion, the clone containing the highest % of CD19^+^ cells was identified and expanded for storage.

### Generation/expansion of CAR-T cells

Frozen peripheral blood mononuclear cells (PBMCs) from adult healthy donors and patients were obtained from Chang Gung Memorial Hospital (IRB approval #201900578A3) and MacKay Memorial Hospital (IRB approval #20MMHIS330e), respectively. PBMCs were extracted using Ficoll-Hypaque gradient separation (Cytiva). Primary human CD3^+^ T cells were isolated from PBMCs by negative selection using EasySep Human T Cell Isolation Kit (StemCell Technologies) or by positive selection using Dynabeads Human T-Expander CD3/CD28 (Thermo Fisher), followed by activation of isolated T cells with Dynabeads (3:1 bead-to-cell ratio) for two days. An iCasp9-CD20/CD19-CAR transposon minicircle carrying CD20 scFv[17] and CD19 scFv[18] and a *qPBase* plasmid were delivered into activated T cells using 4D-Nucleofector device (Lonza) and *Quantum Nufect* Kit (GenomeFrontier Therapeutics) according to the manufacturer’s instructions. Irradiated artificial antigen presenting cells (aAPC), CD19-expressing K562 cells, were added to electroporated T cells at 1:1 ratio on day 3. Cells were cultured in OpTmizer medium (Thermo Fisher) supplemented with *Quantum Booster* (GenomeFrontier Therapeutics) in G-Rex 24- or 6M-well plates (Wilson Wolf) or conventional plates for 10 days. For lentiviral CAR-T production, cryopreserved human PBMCs were thawed and primary human CD3^+^ T cells were isolated from PBMCs by using human Pan T Cell Isolation Kit (Miltenyi Biotec, 130-096-535). Cells were activated and expanded with Dynabeads at 1:1 ratio. After 2 days, cells were transduced with 1 MOI of concentrated virus containing 8 ug/mL Polybrene (Merck, TR-1003-G). Two days later, Dynabeads were removed and the medium (supplemented with IL-7 and IL-15) was changed every 2-3 days until day 10 when CAR-T cells were cryopreserved.

### Cell culture conditions

#### Fed-batch

On day 1, approximately 1×10^5^ nucleofected cells were seeded into a well of 24-well G-Rex containing 3 ml of medium. Fresh medium (2ml) was added on days 3 and 5. On day 7, culture was replaced with fresh medium (2ml).

#### Perfusion

On day 1, nucleofected cells were seeded into conventional 24-well plates at a concentration of 2.5×10^5^ cells/ml. Every 2 to 3 days, the cell numbers were counted, medium was added to adjust the concentration to 2.5×10^5^ cells/ml, and cells were placed back into culture (depending on the volume, up to a 10 ml flask was used to culture these cells).

For both of these culture approaches, cells were harvested on day 10 to be used for subsequent experiments.

### Flow cytometry

CAR expression on T cells was analyzed by co-incubating cells with a Biotin-SP (long spacer) AffiniPure F(ab’)2 Fragment Goat Anti-Mouse IgG antibody (Jackson ImmunoResearch) for 30 min at 4°C and R-phycoerythrin (PE)-conjugated streptavidin (Jackson ImmunoResearch) for 30 min at 4°C. Expression of surface molecules of T cells were determined by labeling T cells with fluorochrome conjugated monoclonal antibodies: CD4-AlexaFlour532 (Thermo Fisher), CD3-Pacific Blue, CD3-PE-Cy5, CD4-AlexaFluor532, CD4-AlexaFlour700, CD8-BV605, CD8-PE-Cy7, CD8-Pacific Blue, CD27-PE-Cy7, CD28-APC, CD45RA-BV421, CD45RO-PE, CD62L-PE-Cy5, CD62L-PE-Dazzle 594, CD95-BV711, CD197-BV510, CD279/PD-1-PE/Cy7, CD366/Tim-3-BV650, CD223/LAG-3-BV711, KLRG-1-BV-421, CD57-PerCP-Cy5.5, CD19-PE, CD56-BV711 (BioLegend), CD45RA-BUV737, and CD95-BUV395 (BD Biosciences). Ghost dye Red 780 fixable viability dye (Cytek), 7AAD or propidium iodide (PI) was used to distinguish live cells from dead cells for data analysis. Mouse blood cell samples were stained with anti-hCD45-APC antibody (BioLegend). For counting the absolute cell number, CountBright™ Absolute Counting Beads (Invitrogen, C36950) were added into FACS samples before acquisition by flow cytometry. Flow cytometric data were acquired on an SA3800 Spectral Analyzer (Sony), a BD FACSCantoTM II (BD Bioscience), or a BD LSRFortessa analyzer (Sony). The graphs were generated using GraphPad Prism or FlowJo software (BD Biosciences or Tree Star).

### Processing mouse blood samples

100μl of blood/mouse was collected in EDTA-containing tubes by submandibular blood collection. After centrifugation at 3000 x g for 15 minutes, plasma (upper layer) was collected for ELISA. The cell pellets/red blood cells (lower layer) were then resuspended with the RBC Lysis Buffer (Biolegend, cat.420301) for 15 minutes at room temperature. Cell pellets were then centrifuged (350 x g for 5 minutes) and washed once with PBS. These cells were used for DNA extraction. For NCI-N87 tumor model experiments, submandibular blood was collected from mice in MiniCollect® tube (K3EDTA, Greiner) and RBC lysed by ACK buffer (Gibco). These cells were used for FACS analysis.

### Enzyme-linked immunosorbent assay (ELISA)

For *in vitro* assay, supernatant was collected 24 hours after CAR-T and Raji-GFP/Luc cell (Creative Biogene, Cat. No: CSC-RR0320) co-culture (E:T=1:1). For *in vivo* assay, mouse plasma samples were collected on days 2, 5, 8, and 14 after T cell injection. IFN-γ (Thermo Fisher), TNF-α (Thermo Fisher), IL-6 (Thermo Fisher), and IL-2 (BioLegend) were quantified by ELISA according to manufacturer’s instructions.

### Genomic DNA extraction and quantitative PCR (qPCR)

Genomic DNA from mouse blood was extracted using a DNeasy Blood & Tissue Kit (Qiagen) following the manufacturer’s instructions. For qPCR analysis, CAR-T cells and Raji-GFP/Luc cells were assessed using the following primer sequences: CAR: fwd-5’-ACGTCGTACTCTTCCCGTCT-3’, rev-5’-GATCACCCTGTACTGCAACCA-3’; Luciferase: fwd-5’-GGACTTGGACACCGGTAAGA-3’, rev-5’-GGTCCACGATGAAGAAGTGC-3’. All qPCR assays were performed using a 7500 fast real-time PCR system (Applied Biosystems). Absolute transgene levels were calculated using the standard curve method. The threshold cycle (CT) value of each target was converted to a concentration using the appropriate standard curve and calculating the % of cells in mouse blood based on the cell copy number.

### *In vitro* cytotoxicity assay

Cytotoxicity was assessed using Celigo image cytometry (Nexcelom). Raji-GFP/Luc target cells were seeded in 96-well culture plates. CAR-T cells were then added at E:T ratios of 5:1, 1:1, and 1:5. At 0, 24, 48, 72, and 96 hours after cell co-culture, Raji-GFP/Luc cells were quantified. The % specific lysis was calculated from each sample using the formula [1-(live fluorescent cell count in target-effector cell co-culture/ live fluorescent cell count in target cell only culture)] x 100.

### Anti-tumor efficacy in mouse tumor models

*In vivo* mouse xenograft model studies were conducted at the Development Center for Biotechnology (Taiwan) using Taiwan Mouse Clinic IACUC (2020-R501-035) approved protocols. Six to eight-week-old male immunodeficient (NOD.Cg-Prkdc^scid^Il2rg^tm1Wjl^/YckNarl) mice (National Laboratory Animal Center, Taiwan) were engrafted with 1.5×10^5^ Raji-GFP/Luc tumor cells intravenously (*i*.*v*.) or with Matrigel (1:1, BD Bioscience) containing 2×10^6^ NCI-N87-Luc cells subcutaneously (*s*.*c*.) on the right flank. For Raji tumor model experiments, mice were infused one week later with 7.5×10^5^ (low), 3×10^6^ (medium), or 1×10^7^ (high) CAR-T cells from donor PBMC-41. After 35 days, 1.5×10^5^ Raji-GFP/Luc tumor cells were injected (*i*.*v*.) to half of the mice in each group. Raji-GFP/Luc tumor cells were quantified using the Xenogen-IVIS Imaging System (Caliper Life Sciences). For NCI-N87 tumor model experiments, mice were injected *i*.*v*. 11 days later with 4×10^6^ control or CAR-T cells. Mice were imaged on an Ami-HT optical imaging system (Spectral Instruments Imaging), following an intraperitoneal (*i*.*p*.) injection of D-Luciferin (BIOSYNTH, L-8220)

### *In vitro* iCasp9 safety assay

Activation of inducible caspase 9 (iCasp9) was performed *in vitro* with 2.5, 5, or 10 nM of dimerizer (AP1903), and CAR-T cell elimination was assessed by flow cytometry 24 hours later.

### ddPCR copy number assays

Genomic DNA (gDNA) was extracted from GF-CART01 cells using a Qiagen DNeasy Blood and Tissue Kit (Qiagen). Minicircle DNA (positive control) was produced by Aldevron. Before ddPCR, the minicircle DNA was digested for 1 min at 37°C with BsrGI. The reaction for ddPCR Copy Number Variation Assay was set up in 20 μl sample volume containing 10 μl of 2X ddPCR Supermix (no dUTP, BioRad), 1 μl of FAM-labeled target primers/probe (BioRad), 1 μl of HEX-labeled reference primers/probe (BioRad), 0.1 μl of BsrGI (Thermo Fisher), and 50 ng of gDNA or minicircle DNA (volume 2 μl). Nuclease-free water was used to adjust the sample to final volume. *RPP30* was used as housekeeping gene. BsrGI directly digests in the ddPCR reaction during setup. Droplets were then generated in a QX200 droplet generator (BioRad) according to manufacturers’ instructions with a final volume of 40 μl. DNA amplification was carried out and droplets were analyzed in a QX200 droplet reader (BioRad). Data was analyzed with QuantaSoft software (BioRad).

### Statistical analysis

Statistical analysis was performed using GraphPad Prism 7 software. Two-tailed unpaired t-tests were used for comparison of two groups. One-way ANOVA was used for comparison of three or more groups in a single condition. *P* values of less than 0.05 were considered statistically significant, \**p* < 0.05, \*\**p* < 0.01, and \*\*\**p* < 0.001.

## RESULTS

### Comparison of fed-batch (G-Rex) and perfusion (conventional)-cultured CAR-T cells

To assess the potential of *qCART*™ system for CAR-T therapy development, we constructed a second-generation bicistronic CAR (GF-CART01) containing binding domains that recognize CD20 and CD19, human 4-1BB and CD3ζ signaling domains, and iCasp9 suicide gene driven by the EF1α promoter **(Figure 1A)**. Following nucleofection of GF-CART01 DNA, we expanded CAR-T cells in G-Rex (fed-batch) and conventional (perfusion) plates. The percentage of CAR^+^ T cells in the fed-batch-cultured (46%) and perfusion-cultured (51%) CAR-T cells were comparable **(Figure 1B)**. However, we observed significantly higher expansion of perfusion-cultured CAR-T cells (836-fold) compared to fed-batch-cultured CAR-T cells (189-fold). Notably, both the perfusion- and fed-batch-cultured CAR-T cells had a high percentage of CD45RA^+^CD62L^+^CD95^+^ T_SCM_ among the CAR^+^ CD4 and CD8 populations. Despite noticeable lower proliferation of fed-batch compared to perfusion-cultured CAR-T cells, we observed enhanced anti-tumor cytotoxicity in fed-batch-cultured CAR-T cells **(Figure 1C)**. As G-Rex will be adopted for good manufacturing practice (GMP) CAR-T manufacturing, all subsequent experiments were conducted in G-Rex.

**Figure 1.**
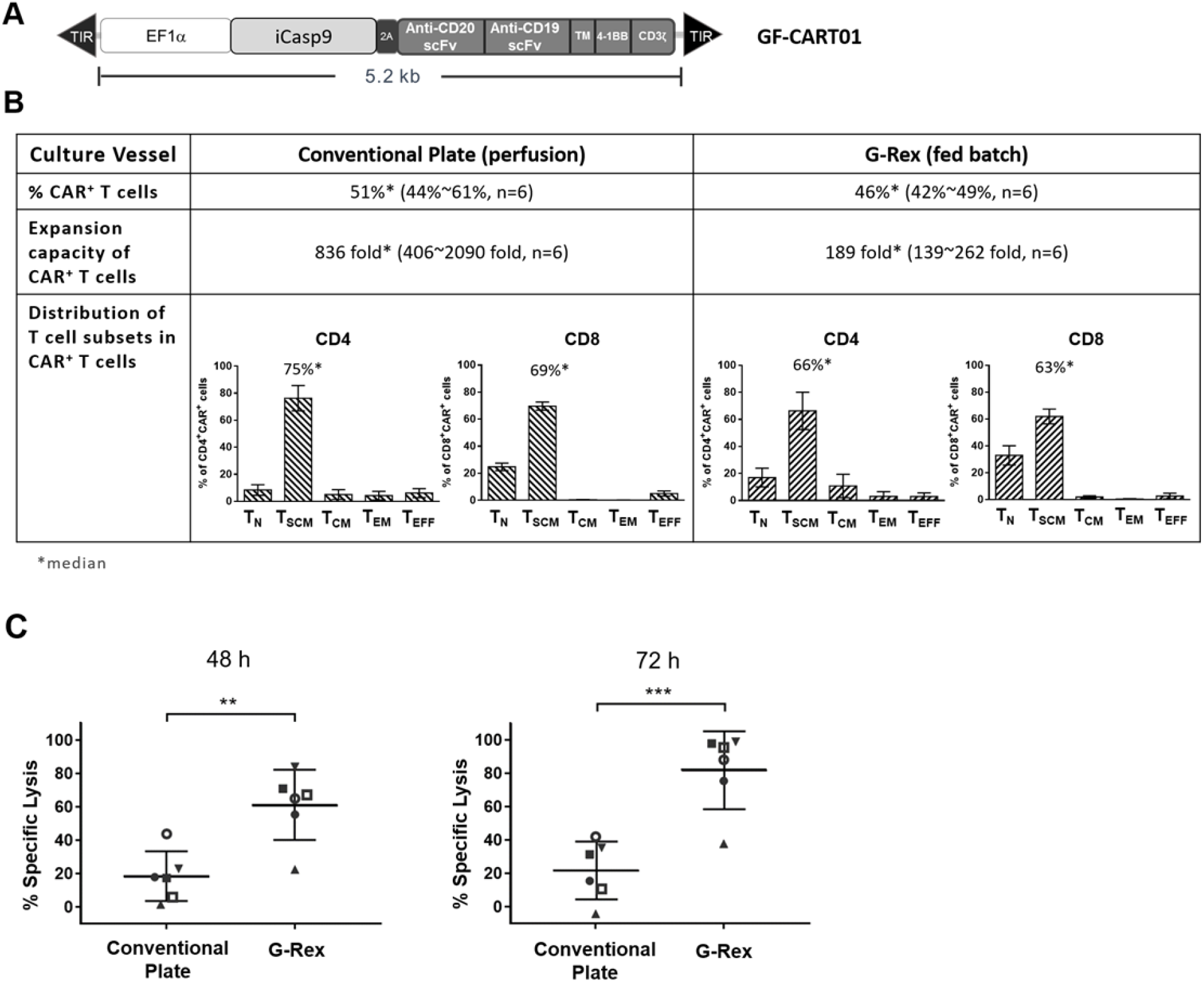
Comparison of perfusion and fed-batch-cultured CAR-T cells. (A) Schematic diagram of the GF-CART01 CAR construct. A T2A sequence enables co-expression of iCasp9 and CD20/CD19-targeted scFvs under the EF1α promoter. (B) Characterization of GF-CART01 cells cultured in conventional plates (perfusion) or G-Rex (fed batch) at day 10 after nucleofection. (C) Cytolytic activities of CAR-T cells after 48 or 72 h of co-culture with Raji-GFP/Luc target cells (E:T ratio of 5:1) were assessed by Celigo image cytometry. (B-C) Data represent mean ± SD in 6 healthy donors, n=6. *p<0.05, \*\**p*<0.01, \*\*\**p*<0.001.

### Safety assessment of GF-CART01 cells

To better manage CAR-T therapy-associated adversities, we included an iCasp9 suicide gene in the GF-CART01 construct. We demonstrated that AP1903 treatment, which activates iCasp9, resulted in a significant reduction in GF-CART01 cell survival **(Supplementary Figure S1A)**. To comply with GMP safety standards for CAR-T manufacturing, we need to ensure that GF-CART01 cells have low integrant (CAR) copy number per cell (<5). We have previously demonstrated that increasing the concentration of DNA for electroporation increased CAR^+^ T cells in a dose-dependent manner (data not shown). Here, we modified CAR-T cells using different amounts of GF-CART01 DNA and determined that the CAR copy number remained low (<5) when 15 μg of total DNA was used for nucleofection **(Supplementary Figure S1B)**. All subsequent experiments were conducted using 15 μg of DNA per 5×10^6^ cells to maximize generation of CAR^+^ T cells while maintaining safety levels of CAR DNA copy numbers.

### GF-CART01 cells are dominated by T_SCM_ cells with high proliferative capacity and low expression of exhaustion and senescence markers

We next examined the proliferative capacity of GF-CART01 cells generated from 9 independent healthy donors and monitored CAR expression on days 1, 8, and 10 following nucleofection. We observed a mean basal level of 21.49% CAR^+^ T cell population on day 1, which subsequently increased and stabilized at 55.7% and 56.75% on days 8 and 10, respectively **(Figure 2A)**. As the presence of transposase (*Quantum PBase*, or *qPBase*) in modified T cells may lead to genotoxicity long-term, we also monitored the *qPBase* levels following nucleofection. While the *qPBase* levels were high on day 1 (47.31%), only a minimal (<0.2%) fraction of *qPBase*^+^ cells was observed on days 8 and 10, suggesting that concerns regarding *qPBase*-induced integrant remobilization should be minimal **(Figure 2B)**. We also observed an average of 178.86-fold GF-CART01 cell expansion **(Figure 2C)**.

**Figure 2.**
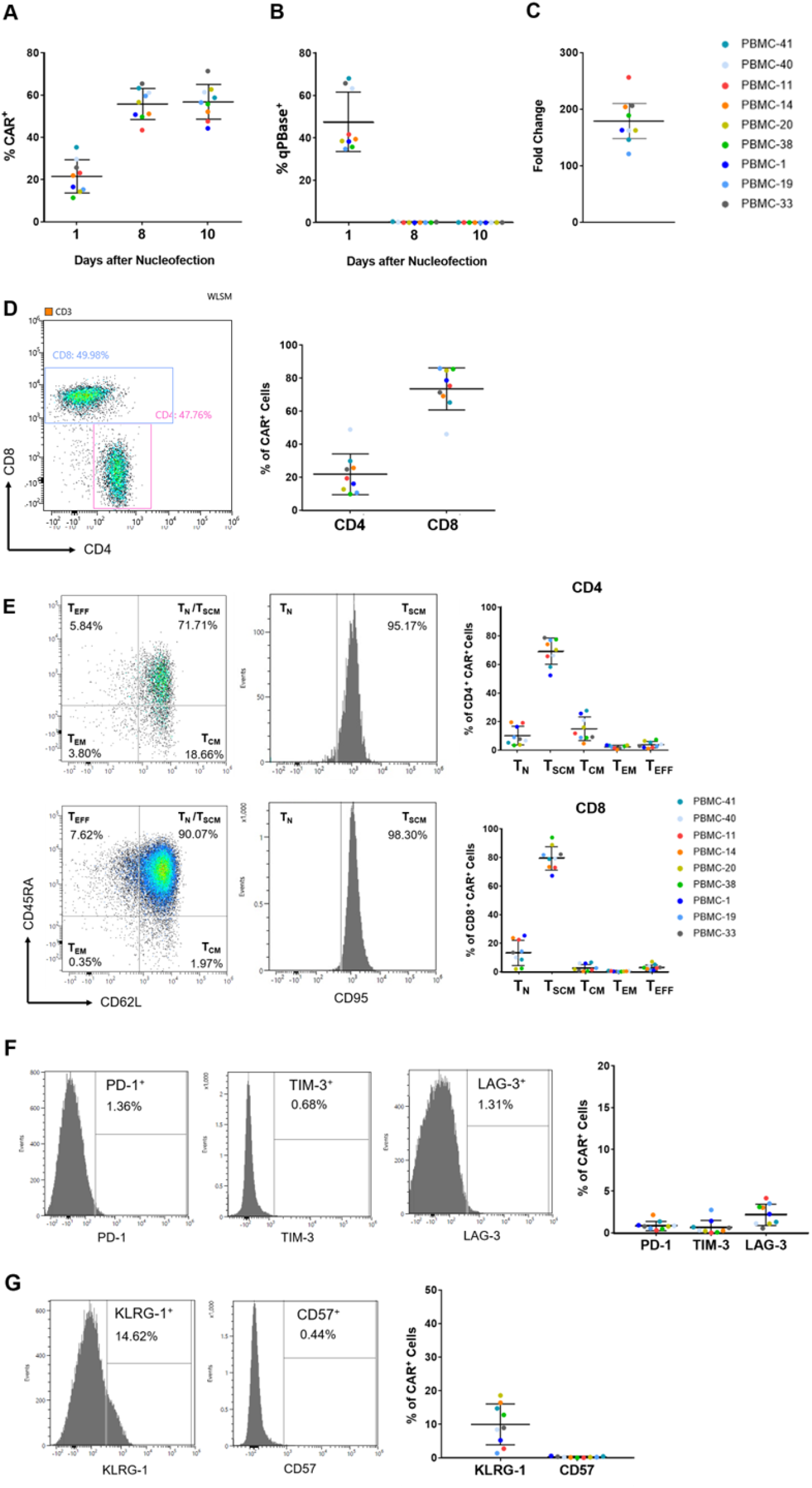
Characteristics of GF-CART01 cells. Percentage of (A) CAR^+^ and (B) transposase^+^ (*qPBase*^+^) cells were analyzed by flow cytometry on days 1, 8, and 10 after nucleofection. (C) Fold change of GF-CART01 cells after 10 days of culture in G-Rex are shown. (D) Percentage of CD4 and CD8 T cells in the CAR^+^ population on day 10 after nucleofection. (E) Distribution of T_N_, T_SCM_, T_CM_, T_EM_, and T_EFF_ subsets in the CD4 (upper panel) and CD8 populations (lower panel). (F) Expression of exhaustion markers PD-1, TIM-3, and LAG-3 in GF-CART01 cells. (G) Expression of senescence markers KLRG-1 and CD57 in GF-CART01 cells. (A-G) Data represent mean ± SD in 9 healthy donors, n=9. Each set of histogram plots shown in (D), (E), (F), and (G) represents one donor.

Among the donors, we consistently observed a higher CD8 (73.58%) compared to CD4 (21.98%) CAR^+^ population **(Figure 2D)**. To assess the composition of GF-CART01 cells, we determined the percentage of CD45RA^+^CD62L^+^CD95^-^ naïve T cell (T_N_), CD45RA^+^CD62L^+^CD95^+^ stem cell memory T cell (T_SCM_), CD45RA^-^CD62L^+^CD95^+^ central memory T cell (T_CM_), CD45RA^-^CD62L^-^CD95^+^ effector memory T cell (T_EM_), and CD45RA^-^CD62L^-^CD95^-^ effector T cell (T_EFF_) subsets in the CD4 and CD8 CAR^+^ populations. We observed high percentage of CAR-T_SCM_ cells in both the CD4 (68.94%) and CD8 (79.84%) populations compared to the other T cell subsets **(Figure 2E)**. Furthermore, these CAR-T cells displayed low exhaustion (PD-1, TIM-3, and LAG-3) markers and the senescence marker CD57 expression **(Figure 2F and G)**. Interestingly, a small but significant population (∼9.92%) expressed the senescence marker KLRG-1.

### GF-CART01 cells display robust cytotoxic function and distinct proliferative responses to CD19- and CD20-expressing target cells

We next tested the anti-tumor function of GF-CART01 cells in two independent donors, PBMC-41 and PBMC-40. Raji-GFP/Luc tumor cells were co-cultured with GF-CART01 cells at effector to target (E:T) ratios of 5:1, 1:1, and 1:5, and cytotoxicity was assessed at 24, 48, 72, and 96 hours after co-culture **(Figure 3A and B)**. While GF-CART01 cells lysed Raji-GFP/Luc tumors in a dose-dependent manner at early time points (24 to 48 h), complete tumor lysis was seen in all E:T ratios at 96 hours. Furthermore, GF-CART01 cells secreted significant levels of pro-inflammatory cytokines IFN-γ, TNF-α, and IL-2 compared to pan-T cells **(Figure 3C-E)**. Collectively, these data suggest that functional GF-CART01 cells can be generated using the *qCART*™ system. Next, we determined how encountering different antigen levels may affect the proliferation capacity of GF-CART01 cells. To address this question, eF450 dye-labeled GF-CART01 cells were co-cultured with K562 cells overexpressing different levels of CD19 or CD20 and proliferation capacity assessed at 48 and 72 hours (see **Supplementary Figure S2** for gating strategy). As shown in **Table 1**, the percentage of proliferated GF-CART01 cells was inversely associated with CD19 expression, but positively associated with CD20 expression, on K562 cells. No such association was observed with CAR^-^ T cells in the corresponding groups and CAR-T cell viability did not appear to be negatively affected by CD19 or CD20 expression (data not shown).

**Figure 3.**
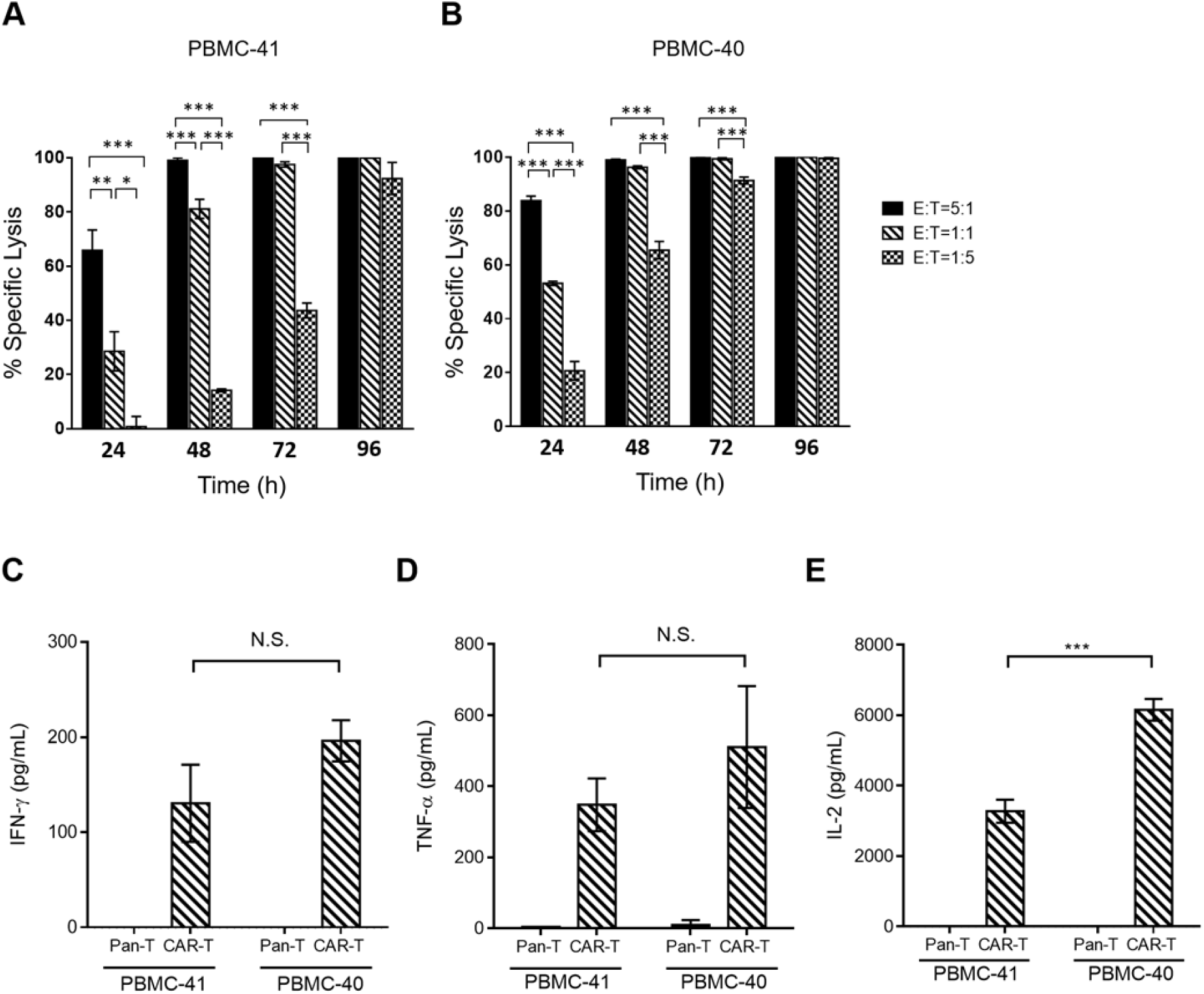
*In vitro* functional analysis of GF-CART01 cells. (A-B) CAR-T cells derived from two healthy donors PBMC-41 and PBMC-40 were assessed for cytotoxicity against Raji-GFP/Luc cells by Celigo image cytometry. Statistical differences were calculated by One-way ANOVA with Tukey multiple comparison, \*\*\**p*<0.001, \*\**p*<0.01, \**p*<0.05. (C) IFN-*γ*, (D) TNF-*α*, and (E) IL-2 secretion by GF-CART01 cells following antigen stimulation was determined by ELISA. Pan-T cells (non-modified cells) served as a control group. Data represent mean ± SD, n=3. Statistical differences are calculated by Student’s t-test, \*\*\**p*<0.001.

**Table 1.** Proliferation capacity of GF-CART01 cells in response to K562 expressing different levels of CD19 or CD20 antigens. Percentages of proliferated live CAR+ and CAR-GF-CART01 cells following 48 h and 72 h co-culture with different levels of CD19- or CD20-overexpressing K562 clones are shown.

|  |  | CAR<sup>+</sup> |  |  |  | CAR<sup>+</sup> |  |  |  |
| --- | --- | --- | --- | --- | --- | --- | --- | --- | --- |
|  |  | % Proliferated |  |  |  | % Proliferated |  |  |  |
|  | Target cell | K562 | K562-CD19 #4-B10 (low) | K562-CD19 #7-B8 (medium) | K562-CD19 #2D-4 (high) | K562 | K562-CD20 #4-2 (low) | K562-CD20 #5-2 (medium) | K562-CD20 #3-2 (high) |
| Time of co-culture | 48 hours | 4.8 ± 1.7 | 28.7 ± 2.4 | 24.9 ± 0.6 | 19.5 ± 1.5 | 7.6 ± 0.8 | 12.5 ± 1.2 | 13.5 ± 0.7 | 11.9 ± 1.3 |
| | 72 hours | 27.6 ± 1.0 | 77.5 ± 1.3 | 69.3 ± 1.9 | 63.9 ± 0.1 | 10.4 ± 0.2 | 60.4 ± 3.0 | 74.7 ± 2.3 | 72.7 ± 2.0 |
|  |  | CAR<sup>-</sup> |  |  |  | CAR<sup>-</sup> |  |  |  |
|  |  | % Proliferated |  |  |  | % Proliferated |  |  |  |
|  | Target cell | K562 | K562-CD19 #4-B10 (low) | K562-CD19 #7-B8 (medium) | K562-CD19 #2D-4 (high) | K562 | K562-CD20 #4-2 (low) | K562-CD20 #5-2 (medium) | K562-CD20 #3-2 (high) |
| Time of co-culture | 48 hours | 2.2 ± 2.0 | 12.6 ± 1.6 | 13.9 ± 0.9 | 11.3 ± 0.3 | 4.7 ± 0.9 | 5.5 ± 2.2 | 5.6 ± 0.7 | 7.2 ± 3.2 |
| | 72 hours | 19.0 ± 1.4 | 44.9 ± 2.6 | 50.3 ± 1.3 | 46.3 ± 0.3 | 6.9 ± 0.8 | 28.1 ± 2.1 | 31.9 ± 0.7 | 25.6 ± 0.5 |

% of CAR<sup>+</sup> cells (stained by anti-mouse IgG Fab') were detected by flow cytometry (SonySA3800) All results were obtained in triplicates

### GF-CART01 cells display enhanced and persistent anti-tumor function *in vivo*

We next evaluated the anti-tumor efficacy of GF-CART01 cells in an *in vivo* xenotransplant immunodeficient mouse model using donor PBMC-41. We injected 1.5×10^5^ Raji-GFP/Luc tumor cells intravenously (*i*.*v*.) followed by 7.5×10^5^ (low), 3.0×10^6^ (medium), or 1×10^7^ (high) GF-CART01 cells (*i*.*v*.) 7 days later as depicted in **Figure 4A**. As assessed by bioluminescence, all mice in the vehicle group succumbed to tumor by day 21 **(Figure 4B-C)**. In contrast, Raji-GFP/Luc tumors were dose-response dependently cleared and all tumors were completely eradicated by day 13 in all three treatment groups and remained tumor-free **(Figure 4B)**. Minimal cytokine production was detected in all three treatment groups on days 2, 8, and 14 **(Supplementary Figure S3A)**. On day 26, no detectable levels of Raji-GFP/Luc tumors were observed in all three treatment groups **(Supplementary Figure S3B)**. Moreover, residual CAR gene was observed in the blood of the medium and high groups, suggesting that many GF-CART01 cells persisted to form memory T cells **(Supplementary Figure S3C)**. We also obtained data from mice injected with non-transfected control Pan-T cells that were cultured under the same conditions as the GF-CART01 cells (**Supplementary Figure S4**). These cells exhibited anti-Raji activity approximately two weeks following injection, which likely resulted from *in vivo* expansion of cells that recognized (and killed) Raji tumor cells via a CAR-independent, TCR-mediated mechanism. However, such a killing effect would not be expected with autologous T cell therapy in patients.

**Figure 4.**
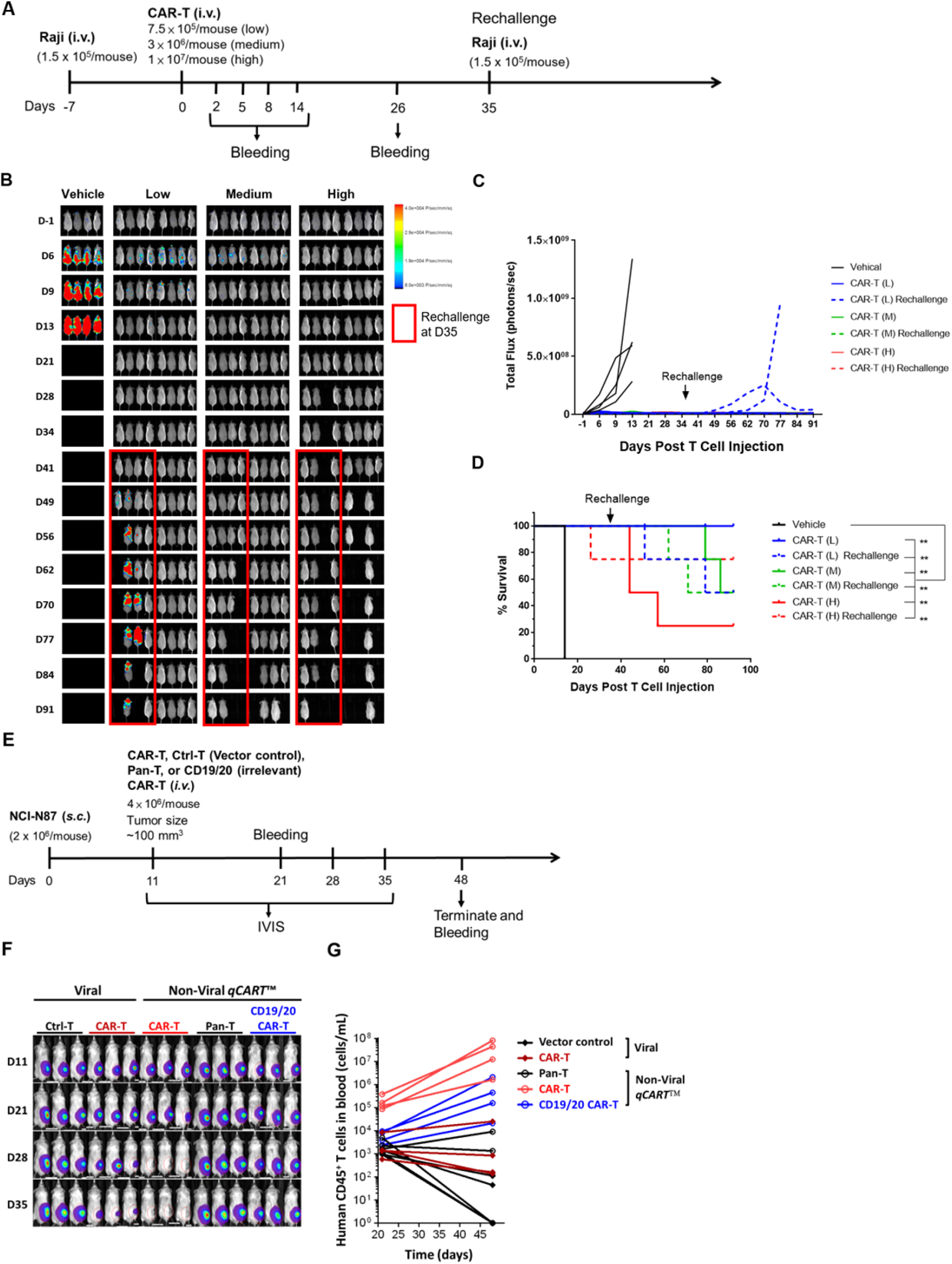
Anti-tumor activity of GF-CART01 cells in a B-cell lymphoma immunodeficient xenograft mouse model. (A) Schematic diagram of *in vivo* Raji xenograft model experimental design. (B-C) Bioluminescent imaging was performed to monitor tumor cell persistence (n= 4/group) and tumor growth were quantified by total flux (photon/s). (D) Kaplan-Meier survival curves for the different treatment groups are shown. (E) Schematic diagram of *in vivo* NCI-N87 xenograft model experimental design. (F) Bioluminescent imaging was performed to monitor tumor cell persistence (n= 3/group). (G) Human T cell counts of blood samples taken from the indicated group of mice at Day 21 and Day 48 post-tumor inoculation.

While residual GF-CART01 cells (as determined by qPCR for presence of CAR gene in genomic DNA of mouse blood samples) can be detected up to day 26, it remained unclear whether these residual cells were memory T cells and retained their anti-tumor function. To address this, we rechallenged half of the mice in each treatment group with 1.5×10^5^ Raji-GFP/Luc tumor cells on day 35. Due to the lack of CAR-T cell persistence, most of the rechallenged mice in the low group eventually succumbed to tumor **(Figure 4B-C)**. In contrast, the rechallenged mice in the medium and high groups remained tumor-free throughout the experiment. Some mice in the medium and high groups did not survive, but this was likely due to graft- versus-host disease (GVHD) as severe ruffling were observed in these mice. More importantly, GF-CART01 cell treatment significantly improved the overall survival of tumor-bearing mice compared to vehicle control **(Figure 4D)**.

To confirm that our non-viral gene engineering approach produces CAR-T cells of high quality and quantity, we next directly compared our non-viral *qCART*™ system with a conventional lentiviral engineering approach. CAR-T cells derived from the same healthy donor PBMC source carrying a CAR gene targeting to a pan-cancer antigen was produced utilizing both approaches. The anti-tumor effects of these CAR-T cells were determined in mice harboring NCI-N87 gastric carcinoma xenografts. The rationale for choosing a single CAR gene targeting to a solid tumor model system was two-fold: First, since the transgene size of GF-CART01used in the Raji tumor model animal experiments was 5.2 Kb (**Figure 4**), CAR-T production using a viral vector system expressing such a sizable transgene would be difficult and may introduce additional variables influencing the results. Second, comparisons carried out in a solid tumor model would be expected to better differentiate between the differences in quality of the CAR-T cells produced using *qCART*™ and the lentivirus systems, since solid tumors are more challenging to be eliminated. We demonstrated that *qCART*™ produced CAR-T cells were superior in performance compared to lentivirally-engineered CAR-T cells, characterized by greater *in vitro* expansion capacity, higher CD8/CD4 ratio, and enrichment of CAR-T_SCM_ cells **(Supplementary Table S1)**. More importantly, in the solid tumor NCI-N87 xenotransplant model **(Figure 4E)**, we observed enhanced antitumor efficacy as well as *in vivo* expansion of *qCART*™-produced CAR-T cells. This was characterized by complete clearance of tumors at Day 28 **(Figure 4F)** in mice treated with *qCART*™-produced CAR-T cells but not those treated with lentivirus-engineered counterparts. Furthermore, between 10 and 37 days following CAR-T injection (CAR-T cells injected at 11 days following tumor inoculation), circulating CAR-T cells robustly proliferated in mice (10 to 100 fold increase in circulating CAR-T cells), while lentivirus-produced CAR-T cells either minimally proliferated or did not proliferate at all **(Figure 4G)**. This difference may reflect the above-mentioned higher CD8/CD4 ratio and percentage of T_N_/T_SCM_ population present in both the CD4^+^ and CD8^+^ CAR-T populations of *qCART*™-produced CAR-T cells **(Supplementary Table S1)**.

### Functional GF-CART01 T_SCM_ cells can be generated in sufficient numbers from cancer patients

We next assessed whether sufficient high-quality GF-CART01 cells can be generated for clinical applications. In our previous experiments thus far, we expanded all GF-CART01 cells with CD19-expressing aAPC. To streamline the manufacturing process, we tested whether we can still generate highly potent patient-derived GF-CART01 T_SCM_ cells in the absence of aAPC selective expansion. We isolated PBMCs from 6 healthy donors, 3 diffuse large B-cell lymphoma (DLBCL) patients, 3 chronic lymphocytic leukemia (CLL) patients, one Hodgkin lymphoma (HL) patient, and one multiple myeloma (MM) patient to generate GF-CART01 cells. **Figure 5** and **Table 2** summarize the characteristics of the healthy donor- and patient-derived CAR-T cells.

**Figure 5.**
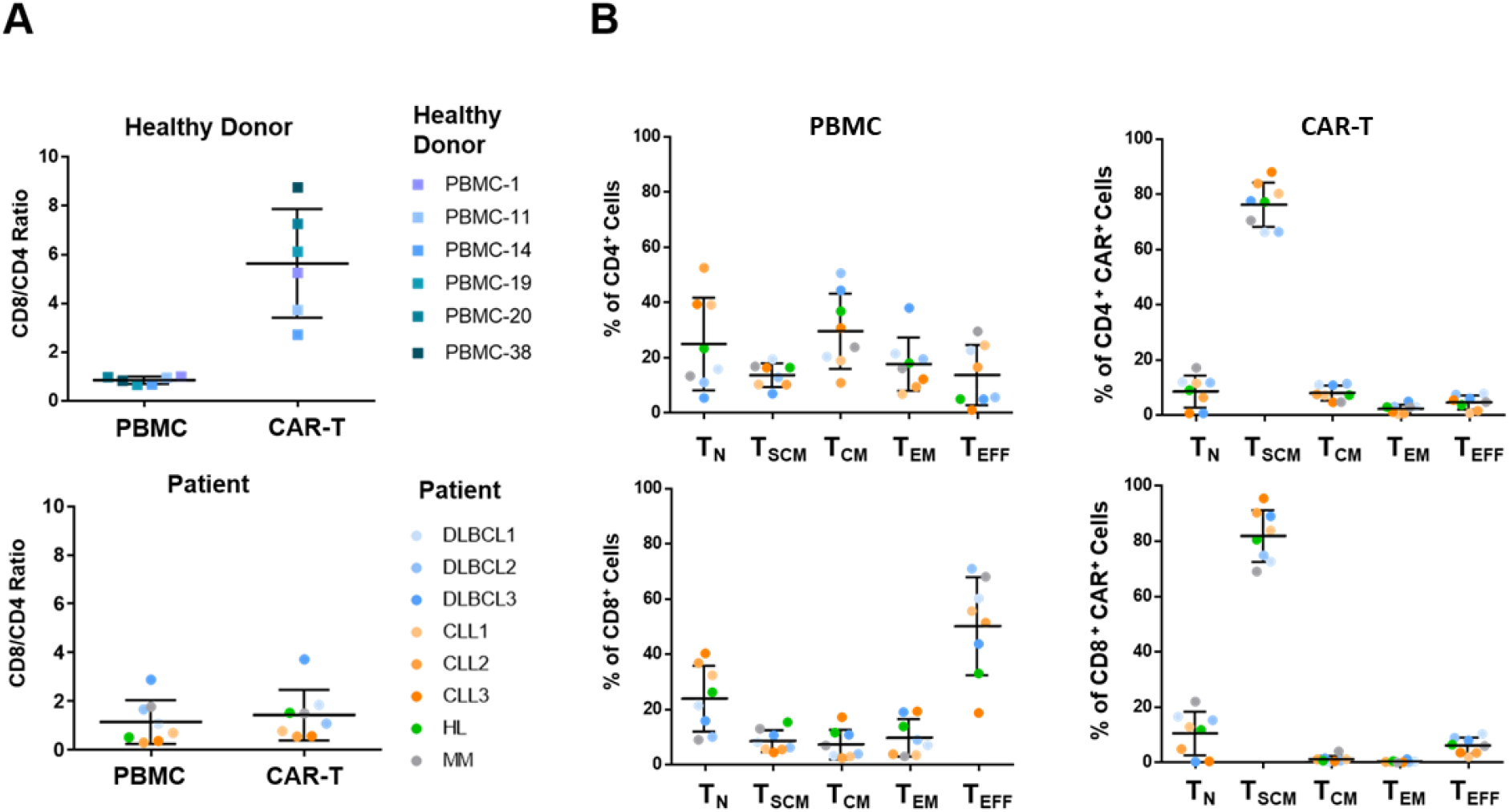
Assessment of cancer-patient-derived GF-CART01 cells. T cells derived from healthy donors and cancer patients were nucleofected with GF-CART01. (A) CD8/CD4 ratios were assessed before (PBMC) and after (CAR-T) nucleofection. (B) Distribution of T_N_, T_SCM_, T_CM_, T_EM_, and T_EFF_ subsets in CD4^+^ (upper panel) and CD8^+^ (lower panel) CAR-T cell populations. DLBCL, diffuse large B-cell lymphoma; CLL, chronic lymphocytic leukemia; HL, Hodgkin’s lymphoma; MM, multiple myeloma.

**Table 2.**
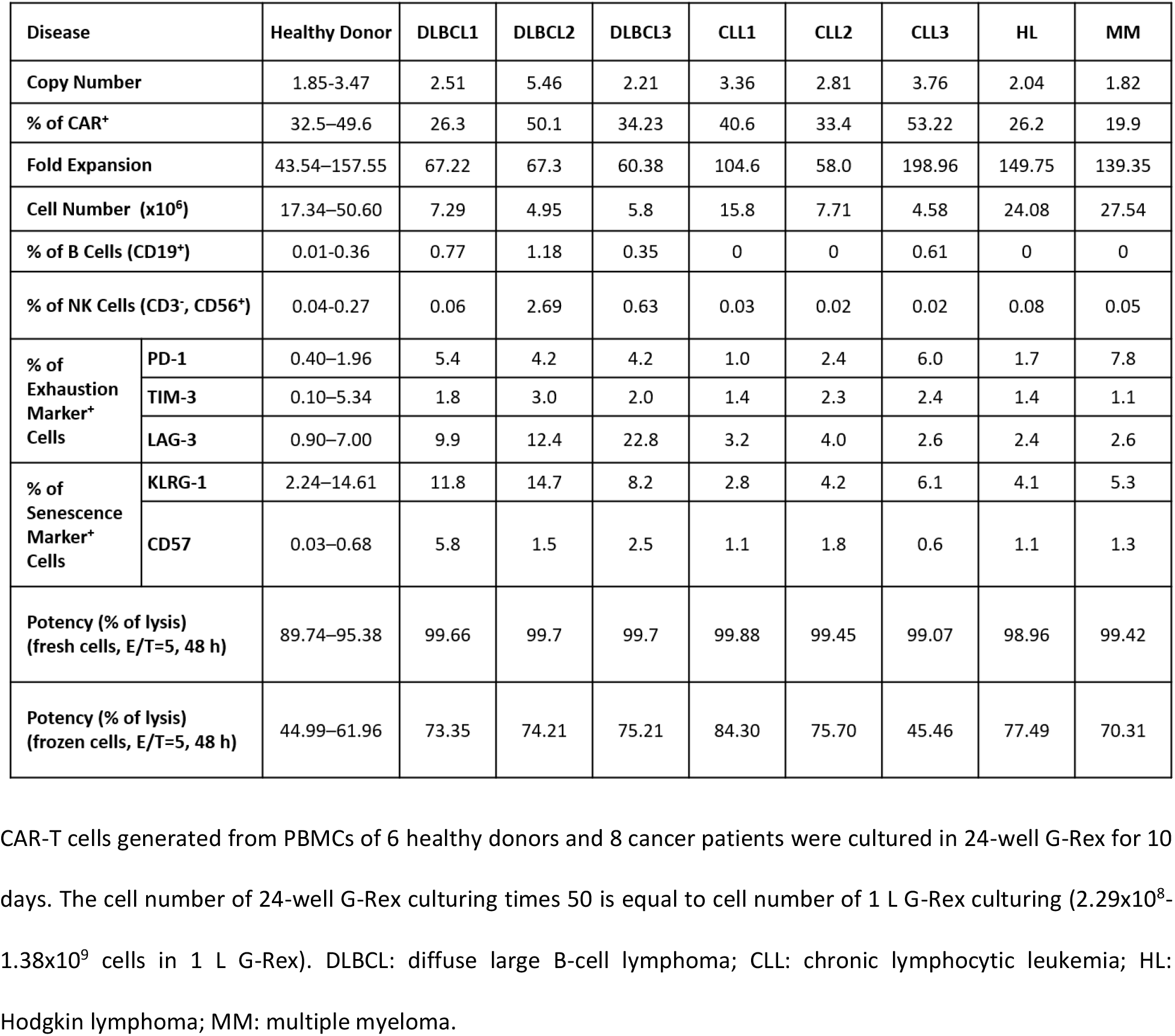
Comparison of GF-CART01 cells generated from healthy donors and cancer patients.

Over the course of 10 days, we expanded the CAR-T cells to 4.58×10^6^–2.76×10^7^ cells in a 24-well G-Rex culture, which would be equivalent to clinically relevant number of 2.29×10^8^–1.38×10^9^ cells in a 1 L G-Rex culture vessel. The CAR expression (19.9–53.22%) and the fold expansion (58.0–198.96 folds) of the patient cells were comparable to those from healthy donors (32.5–49.6% and 43.54–157.55 folds) **(Table 2)**. The CAR-T cell purity was confirmed by the low percentage of CD19^+^ B cells and CD3^-^CD56^+^ NK cells in both the healthy donors (0.01-0.36% B; 0.04-0.27% NK) and patient cells (0-1.18% B; 0.02-2.69% NK) **(Table 2)**. The proportion of CD8 and CD4 were balanced in both the healthy donors (CD8/CD4=0.85) and the patients (CD8/CD4=1.17) prior to nucleofection **(Figure 5A)**. After nucleofection, the CAR-T cells from healthy donors had higher CD8/CD4 ratio (5.64). However, the increase in the proportion of CAR^+^ CD8 T cells was only observed in some patients **(Figure 5A)**. Notably, compared to pre-nucleofection (PBMC) levels, the percentages of T_SCM_ subset at 10 days post-nucleofection increased in all patients among the CD4 (from 13.75% to 76.25%) and CD8 (from 7.17% to 81.90%) populations, confirming the unique quality of *qCART*™ system in supporting T_SCM_ differentiation **(Figure 5B**, see **Supplementary Figure S5** for gating strategy). To further confirm T_SCM_ status, we determined the expression of additional surface antigen biomarkers whose expression have previously been reported to be associated with T_SCM_ and compared their expression among the T_SCM_ (CD45RA^+^CD62L^+^CD95^+^) population of PBMC T cells and CAR-T cells derived from three patients and healthy volunteers (**Supplementary Figure S6**). Consistent with data presented in **Figure 2E** and **Figure 5B**, after engineering and expansion, both the patients’ and healthy volunteers’ (CD4 and CD8) CAR-T cells were highly enriched for T_SCM_ (**Supplementary Figure S6A**). High percentage of these T_SCM_ cells also co-expressed CD45RO, CCR7, CD27 and CD28, confirming these cells’ T_SCM_ status (**Supplementary Figure S6C-S6D**). Notably, the CD45RO^dim^ population in these T_SCM_ cells may represent cells that were in transition into a more differentiated state. Furthermore, we also found that the T_N_/T_SCM_ population of (particularly healthy donor) PBMCs contained only low percentage of CD95^+^ cells (**Supplementary Figure S6B**), suggesting that the T_N_/T_SCM_ population of PBMCs were dominated by naïve T cells. The GF-CART01 cells were also assessed for expression of exhaustion markers PD-1 and TIM-3 and senescence marker CD57, and only a low percentage of cells were found to express these markers **(Table 2)**. In addition, the percentage of cells expressing the exhaustion marker LAG-3 was low in CLL, HL, and MM patients (2.4-4%) with a slightly elevated percentage of cells expressing these markers among the DLBCL patients (9.9-22.8%). While KLRG-1 expression (∼7.25%) was detectable in GF-CART01 patient cells, it was significantly lower than KLRG-1 expression prior to nucleofection (∼21.27%; p=0.0087) **(Supplementary Figure S7A)**. The decrease in KLRG-1 expression was especially prominent in the CD8 population (52.50% vs. 14.59%; p=0.0004) **(Supplementary Figure S7B)**. A 48-hour cytotoxicity assay further confirmed the functionality of the GF-CART01 cells **(Table 2)**. Together, our data suggest that the *qCART*™ system is a promising technology for generating functional multiplex armored CAR-T_SCM_ cells for cancer immunotherapy application. GF-CART01 therapy is a safe, potent, and cost-effective treatment for B-cell malignancies.

## DISCUSSION/ CONCLUSION

*qPB* is currently the most efficient transposon for gene integration, and GF-CART01 cell therapy was the first CAR-T therapy developed using the *qPB*-based *qCART*™ system.[15,16] The *qCART*™ system can easily produce CAR-T cells with transgene size >7.6 kb, which is not achievable by conventional lentiviral/retroviral CAR-T engineering.[16] The limited cargo capacity of lentiviral/retroviral vector may account for the lack of current CAR-T clinical trials targeting B-cell malignancies with both dual-targeting and iCasp9 safety switch gene (5.2 kb transgene size) in the CAR design. In this report, we presented evidence on the feasibility of manufacturing highly potent iCasp9-regulatable CD20/CD19-targeted CAR-T_SCM_ cells using the virus-free *qCART*™ system for clinical application. We demonstrated the purity, potency, and consistency profiles of CAR-T cells manufactured using the *qCART*™ system.

Maintaining sterility and controlling mycoplasma and endotoxin contamination in the CAR-T product are important safety concerns. G-Rex is a GMP-compliant medical device used for cell expansion.[19,20] The minimal manual operation of this device minimizes contamination risks. Importantly, manufacturing GF-CART01 cells with G-Rex can be achieved without compromising CAR-T efficacy. Consistent with our previous study, we observed low risk of genotoxicity in GF-CART01 cells, given the safe integration profile, low enhancer activity, and low residual *qPBase*.[16] Acute toxicities in CAR-T therapy-treated patients, such as CRS and ICANS, have been major concerns.[21–23] Furthermore, a rare case of CAR^+^ CD4 T cell lymphoma has been reported recently in patients treated with CD19 CAR-T cells.[24,25] To safely manage toxicities, CAR-T cells can be eradicated by targeting CAR-T cells (e.g. anti-EGFRt antibody) via antibody-dependent cellular cytotoxicity (ADCC) mechanism or using a small molecule that activates a suicide gene introduced in the CAR design (e.g. iCasp9).[21] The ADCC approach has slower onset of killing compared to the latter. We included an iCasp9 safety switch in our GF-CART01 design to mitigate the adverse effects of CRS and ICANS. Indeed, Foster et al (2021) has recently demonstrated that acute ICANS can be effectively controlled by treating CAR-T therapy patients with the iCasp9-activating agent, rimiducid (AP1903).[26]

Manufacturing clinically relevant number of a pure high-quality CAR-T product is the first critical step towards successful CAR-T therapy. One approach to expanding the CAR^+^ T cell population is by culturing with aAPC, but this often leads to a more differentiated and short-lived T_EFF_ or exhaustive CAR-T phenotype.[27] Another approach is to include markers within the construct to enrich the CAR^+^ T cell population either via drug selection or antibody-mediated cell isolation during or after CAR-T *ex vivo* expansion, respectively. However, such approaches increase the payload size and introduce therapeutically irrelevant genes into the genome.[4,22,23] While we also used aAPC in our initial experiments, we could generate sufficient quantity of patient-derived GF-CART01 cells without aAPC-induced enrichment by addition of *Quantum Booster*™, a serum-free supplement which contains cellular proteins and a set of cytokines with defined concentrations. These components of *Quantum Booster*™ act to promote proliferation of the CAR-T cells while maintaining stemness as well as minimal levels of senescence and exhaustion.[16] Notably, we have shown that *Quantum Booster*™ highly enriches CD4 T_SCM_, which may be critical in maintaining CD8^+^ CAR-T cells at the T_SCM_ differentiation stage while being robustly expanded. Our recent study further confirmed that *qCART*™-manufactured CAR-T cells may be more effective in tumor control without aAPC enrichment.[16] Here, we demonstrated that 2.29×10^8^– 1.38×10^9^ T cells could be generated in a 1 L-G-Rex within a short time-frame of 10 days using cancer patient-derived T cells. Even at a low CAR^+^ percentage of 20%, we could still produce sufficient number of CAR^+^ CAR-T cells (4.58×10^6^–2.76×10^7^) for treatment, and we can easily enrich CAR^+^ T cells with aAPC should the need arise.

The high potency of GF-CART01 cells can be attributed to the balanced CD8/CD4 CAR-T ratio and the high enrichment of T_SCM_ in the engineered cells. It has been reported that a balanced CD8/CD4 CAR-T ratio resulted in high remission rates.[28,29] To achieve this, others have engineered CD4 and CD8 CAR-T cells separately before adoptive transfer.[28] While *qCART*™ cells derived from healthy donors generally favored CD8 enrichment, cancer patient-derived *qCART*™ cells have a balanced CD8/CD4 CAR-T ratio (**Figure 5)**. Thus, the *qCART*™ system naturally achieves the desired CD8/CD4 balance without the need to engineer CD4 and CD8 CAR-T cells separately as seen in SB100X-modified CAR-T clinical trials.[30]

Our observation that, in contrast to a positive dose-dependent relationship between CD20 target cell expression level and proliferation capacity of GF-CART01 cells, there appeared to be an inverse association between CD19 target cell expression level and CAR-T proliferation when encountering K562 cells overexpressing different levels of CD19. However, potential toning of GF-CART01 cells would be unlikely since we found CD19 expression level of Raji cell line was much lower than the K562 clone which expressed the lowest level of CD19. In other words, since tumor’s endogenous CD19 expression level was generally lower than the level that can inhibit CAR-T proliferation, we do not expect to see a decrease in CAR-T proliferation similar to that observed with the K562 CD19 overexpression clone which expressed high levels of CD19.

T_SCM_ are a class of immunological memory critical for successful adoptive T cell therapies.[31–35] The *qCART*™ system is unique in its ability to enrich T_SCM_ population without compromising the cell quality, expansion, and production time of CAR-T cell product.[16] One approach to harness the potency of less-differentiated T cells such as T_SCM_ is to shorten the *ex vivo* expansion time of CAR-T product. *Ghassemi et al*. (2022) demonstrated that shortening the lentiviral CAR-T manufacturing process to 24-hours resulted in a higher percentage of CAR-T cells with a memory phenotype.[36] Though these CAR-T cells displayed stronger *in vivo* anti-tumor activity than their conventional counterpart, successful clinical translation of this process remains to be demonstrated. Another approach is to culture CAR-T cells with specific cytokines or chemical reagents.[34,35,37–39] Additionally, CAR-T cell production from defined T cell subsets has been shown to be beneficial in enriching the T_SCM_ population.[40] However, none of these approaches enrich T_SCM_ population to a high level seen in our *qCART*™-modified T cells. While T_SCM_ cells make up a small percentage of T cells derived from cancer patients, genetically modifying these cells using the *qCART*™ system resulted in enrichment of T_SCM_ to as high as >80% in both the CD4 and CD8 CAR^+^ T cell populations.

In conjunction with the ability of T_SCM_ to self-renew and persist long-term, we also observed low expression of exhaustion (PD-1, TIM-3, and LAG-3) and senescence (CD57) biomarkers in our GF-CART01 cells **(Figure 2F-G; Table 2)**. KLRG-1 is a senescence marker. While we observed a small but significant population of KLRG-1^+^ cells in GF-CART01 cells, we also observed a significantly lower percentage of KLRG-1^+^ cells after CAR-T production compared to freshly isolated PBMCs **(Supplementary Figure S2)**. Herndler-Brandstetter *et al*. (2018) recently described a class of memory T cell population developed from KLRG-1^+^ effector CD8 T cells that have lost KLRG-1 expression.[31] These so-called “exKLRG-1 memory cells” have been demonstrated to inhibit tumor growth more efficiently compared to KLRG-1^+^ cells in an OT-I melanoma mouse model. T_SCM_ cells generally lack KLRG-1 expression, and KLRG-1^+^ T_SCM_ cells have been associated with cancer patients undergoing relapse.[32,33] Given the significant decrease in KLRG-1 expression in GF-CART01 cells, it is possible that a significant proportion of CAR-T_SCM_cells in our model were actually exKLRG-1 memory cells. Thus, one possible mechanism that made it possible for *qCART*™ system to generate highly potent CAR-T_SCM_ cells may be through the conversion of KLRG-1^+^ effector CD8 T cells to exKLRG-1 memory cells.

One of the major challenges of CAR-T cell manufacturing is the variation and inconsistency of CAR-T cell products.[34] In one study of ten CLL and ALL patients, CAR-T cells expanded anywhere between 23.6 and 385-folds with retroviral CAR transduction of 4-70%.[35] Lentiviral transduction of 5.5-45.3% was reported by other studies of CLL and ALL patients.[36,37] In our GF-CART01 study, we observed tighter range of CAR^+^ percentage (19.9-53.22%) and expansion (58.0-198.96 folds) among the eight cancer patients **(Table 2)**. Furthermore, these low levels of variation were comparable to the CAR^+^ percentage (32.5-49.6%) and expansion (43.54-157.55 folds) among the healthy donors in our study, respectively. More importantly, GF-CART01 cells derived from both healthy donors and patients can effectively eradicate tumor cells.

In this study, we directly compared between the performance and anti-tumor efficacy of *qCART*™- and lentivirally-engineered CAR-T cells in a solid tumor xenograft model. The results clearly demonstrated superior performance, anti-tumor efficacy, and *in vivo* expansion of *qCART*™-produced CAR-T cells over lentivirally-produced CAR-T cells (**Figure 4E-4G**). These observations suggest that *qCART*™-produced CAR-T cells, including GF-CART01 cells, are indeed highly potent.

In summary, the *qCART*™ system is an elegant system for the manufacturing of CAR-T products. In this study, we demonstrated that the *qCART*™ system can be used to produce patient-derived GF-CART01 cells with all the desired attributes of CAR-T products to ensure therapeutic success, including high CAR-T_SCM,_ balanced CD8/CD4 ratio, low exhaustion and senescence marker expressions, high expansion capacity, and excellent anti-tumor efficacy. We believe that the simplicity and robustness of manufacturing multiplex CAR-T cells with enhanced product consistency using the *qCART*™ system will not only significantly enhance the accessibility of CAR-T therapy but also unlock the full potential of armored CAR-T therapy for the treatment of solid tumors in the future.

## List of Abbreviations

aAPC: artificial antigen presenting cells
CAR: chimeric antigen receptor
CLL: chronic lymphocytic leukemia
CRS: cytokine release syndrome
CT: threshold cycle
DLBCL: diffuse large B-cell lymphoma
EFS: event-free survival
GF-CART01: patient-derived bicistronic CD20/CD19-targeted iCasp9 regulatable CAR-T cells
GMP: good manufacturing practice
GVHD: graft-versus-host disease
HL: Hodgkin lymphoma
ICANS: immune effector cell-associated neurotoxicity syndrome
iCasp9: inducible caspase 9
i.v.: intravenously
MM: multiple myeloma
NHL: Non-Hodgkin lymphoma
PBMCs: peripheral blood mononuclear cells
PE: R-phycoerythrin
PFS: progression-free survival
*qCART*: *Quantum CART*
*qPB*: Quantum pBac
T_CM_: central memory T cell
T_EFF_: effector T cell
T_N_: naïve T cell
T_SCM_: T stem cell memory

## Acknowledgements

The authors thank Ms. Jin-hwa Yang for her assistance with IRB preparation and approval process. The authors also thank Dr. Pei-Yi Tsai for assistance with the animal experiments.

