## Supplementary Materials for "Manufacturing CD20/CD19-targeted iCasp9 regulatable CAR-T_SCM_ cells using *qCART*, the *Quantum pBac*-based CAR-T system"

**Running Title: Manufacturing CAR-$T_{SCM}$ cells using *qCART***

Peter S. Chang[1,*], Yi-Chun Chen[1,*], Wei-Kai Hua[1,*] , Jeff C. Hsu[1], Jui-Cheng Tsai[1], Yi-Wun Huang[1], Yi-Hsin Kao[1], Pei-Hua Wu[1], Yi-Fang Chang[2,3,4], Ming-Chih Chang[2], Yu-Cheng Chang[2,3], Shiou-Ling Jian[5], Jiann-Shiun Lai[5], Ming-Tain Lai[5], Wei-Cheng Yang[6], Chia-Ning Shen[6,7], Kuo-Lan Karen Wen[1,†], Sareina Chiung-Yuan Wu[1,†]

[1]GenomeFrontier Therapeutics, Inc. Taipei City, Taiwan (R.O.C.)

[2]Division of Hematology and Oncology, Department of Internal Medicine, Mackay Memorial Hospital, Taipei, Taiwan (R.O.C.)

[3]Department of Medical Research, Laboratory of Good Clinical Research Center, Mackay Memorial Hospital, Tamsui District, New Taipei City, Taiwan (R.O.C.)

[4]Department of Medicine, Mackay Medical College, New Taipei City, Taiwan (R.O.C.)

[5]OBI Pharma, Inc.

[6]Biomedical Translation Research Center, Academia Sinica, Taipei, Taiwan (R.O.C.)

[7]Genomics Research Center, Academia Sinica, Taipei, Taiwan (R.O.C.)

[*]These authors contributed equally to this work and their authorship order is interchangeable

[†]To whom correspondence may be addressed. Sareina Chiung-Yuan Wu, Kuo-Lan Karen Wen; 18F-1, No. 3, Park St., Nangang Dist., Taipei City 11503, Taiwan (R.O.C.); +886-2-26558766;

**This supplementary information file contains:**

(1) Supplementary Figure Legends; and

(2) Figures S1-S7 and Table S1

**Supplementary Figure Legends**

**Supplementary Figure S1. Safety assessment of GF-CART01 cells.** (A). Elimination of GF-CART01 cells after iCasp9 activation. Percentage of $CAR^{+}$ T cells were assessed by flow cytometry at 24h after AP1903 treatment. Data represent mean ± SD mean, n=3 (One-way ANOVA with Tukey multiple comparison). *p<0.05, ***p*<0.01, ****p*<0.001. (B) Average CAR copy numbers were assessed in GF-CART01 cell products following electroporation with different amounts of DNA using primers against the scFv CAR region by ddPCR, n=2 or 3.

**Supplementary Figure S2. Effect of CD19 and CD20 expression level on proliferation capacity of GF-CART01 cells.** (A) Flow cytometric analysis showing the mean fluorescence intensity (MFI) levels of CD19 and CD20 staining on Raji, K562 and low, medium and high level CD19- or CD20-overexpressing K562 clones. (B) A representative set of histogram plots showing gating strategy to determine proliferated CAR-T cells.

**Supplementary Figure S3. Anti-tumor activity of GF-CART01 cells in a B-cell lymphoma immunodeficient xenograft mouse model.** (A) Plasmas from mice were collected on days 2, 5, 8, and 14 after CAR-T cell administration and analyzed for IFN-$\gamma$, TNF-$\alpha$, IL-2, and IL-6 by ELISA. Copy number of (B) luciferase and (C) CAR were determined in mouse blood samples on day 26 after CAR-T cell administration to monitor the presence of $CAR^{+}$ T cells and Raji-GFP/Luc cells, respectively, in the mouse circulation (n=8 and n=7 for

CAR-T(M) and CAR-T(H)). Dotted lines represent the detection limits of qPCR. In (B) and (C), dotted lines represent the detection limit.

**Supplementary Figure S4. Anti-tumor activity of GF-CART01 cells in a B-cell lymphoma immunodeficient xenograft mouse model.** (A) Schematic diagram of *in vivo* experimental design. (B) Bioluminescent imaging was performed to monitor tumor cell persistence (n= 4).

**Supplementary Figure S5. Gating strategy for $T_{SCM}$ cells.** CD4 or CD8 $T_{SCM}$ phenotyping and quantification strategy for analysis of (A) PBMCs were performed by sequential gating on (1) lymphocytes without debris, (2) $PI^-$ or $7AAD^-$ (live) cells, (3) $CD3^+$ cells, (4) either $CD4^+$ or $CD8^+$ cells, (5) $CD45RA^+$ and $CD62L^+$ cells, and (6) $CD95^+$ cells. (B) CAR-T cells were analyzed as in (A), except in step (2) $PI^-$ or $7AAD^-$ (live) and $CAR^+$ cells are gated on. Shown are representative data plots of patient DLBCL3.

**Supplementary Figure S6. Characteristics of GF-CART01 cells (additional samples).** PBMCs and CAR-T cells derived from healthy donors and cancer patients were nucleofected with GF-CART01. (A) Distribution of $T_N$, $T_{SCM}$, $T_{CM}$, $T_{EM}$, and $T_{EFF}$ subsets in the CD8 (upper panels) and CD4 populations (lower panels). (B) Percent of $CD95^+$ cells in the CD8 (upper panels) and CD4 $T_N/T_{SCM}$ populations (lower panels). (C) A representative data set showing percentage of $CD45RO^+$, $CCD7^+$, $CD27^+$ and $CD28^+$ cells in the $CAR^+CD8^+T_{SCM}$ (upper panels) and $CAR^+CD4^+T_{SCM}$ populations (lower panels). (D) Percentage of $CD45RO^+$, $CCD7^+$, $CD27^+$ and $CD28^+$ cells in the $T_{SCM}$ cells obtained from healthy donors (left panel) and patients (right panel).

**Supplementary Figure S7. Comparison of exhaustion and senescence before and after CAR nucleofection.** (A) Expression of exhaustion (PD-1, TIM-3, and LAG-3) and senescence (KLRG-1 and CD57) markers in cancer patients from **Table 2** before (PBMC) and after (CAR-T) nucleofection. (B) Expression of KLRG-1 within the CD8 population of the cancer patients.

**Supplementary Table S1. Performance of CAR-T cells produced using lentiviral and non-viral *qCART*™ cell production.**

**Figure S1**

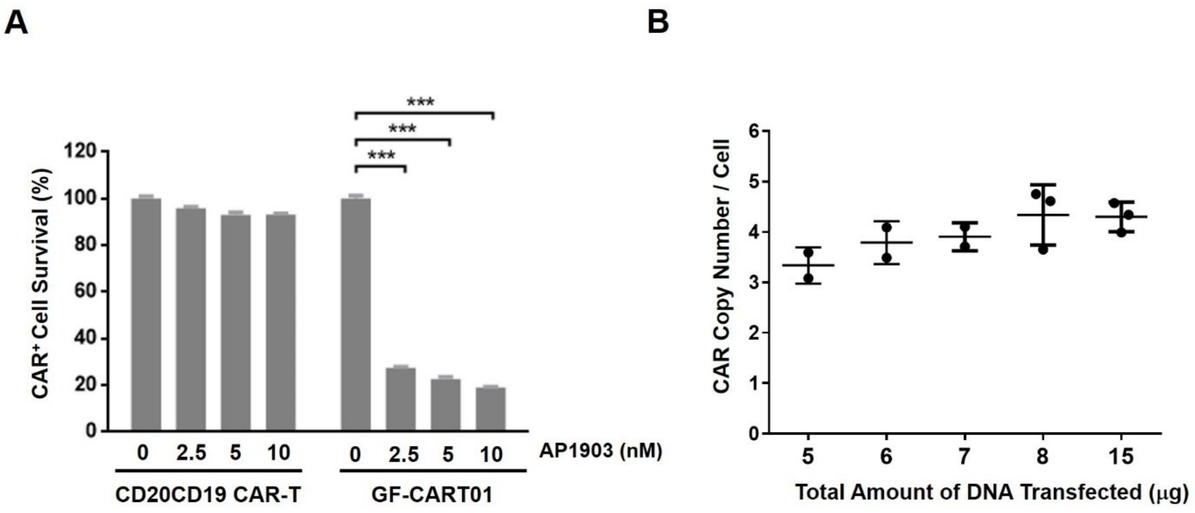
A
B
***
***
***
120
100
80
60
40
20
0
CAR⁺ Cell Survival (%)
0
2.5
5
10
0
2.5
5
10
AP1903 (nM)
CD20CD19 CAR-T
GF-CART01
6
5
4
3
2
1
0
CAR Copy Number / Cell
5
6
7
8
15
Total Amount of DNA Transfected (μg)

**Figure S2**

**A**

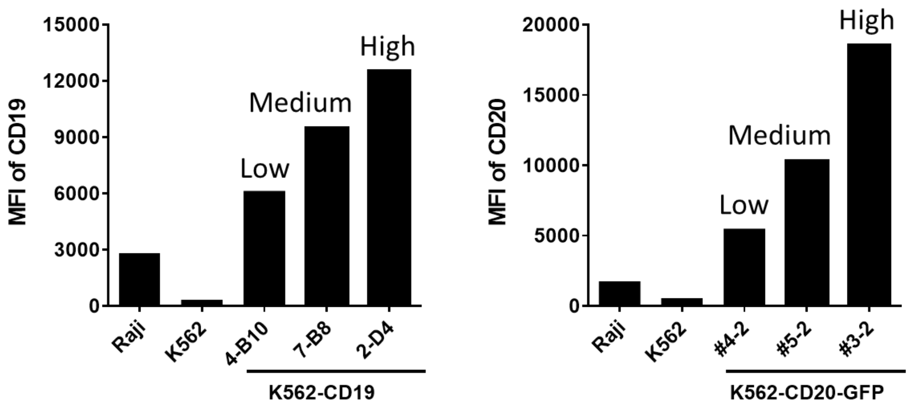

MFI of CD19
15000
12000
9000
6000
3000
0
High
Medium
Low
Raji
K562
4-B10
7-B8
2-D4
K562-CD19
MFI of CD20
20000
15000
10000
5000
0
High
Medium
Low
Raji
K562
#4-2
#5-2
#3-2
K562-CD20-GFP


**B**

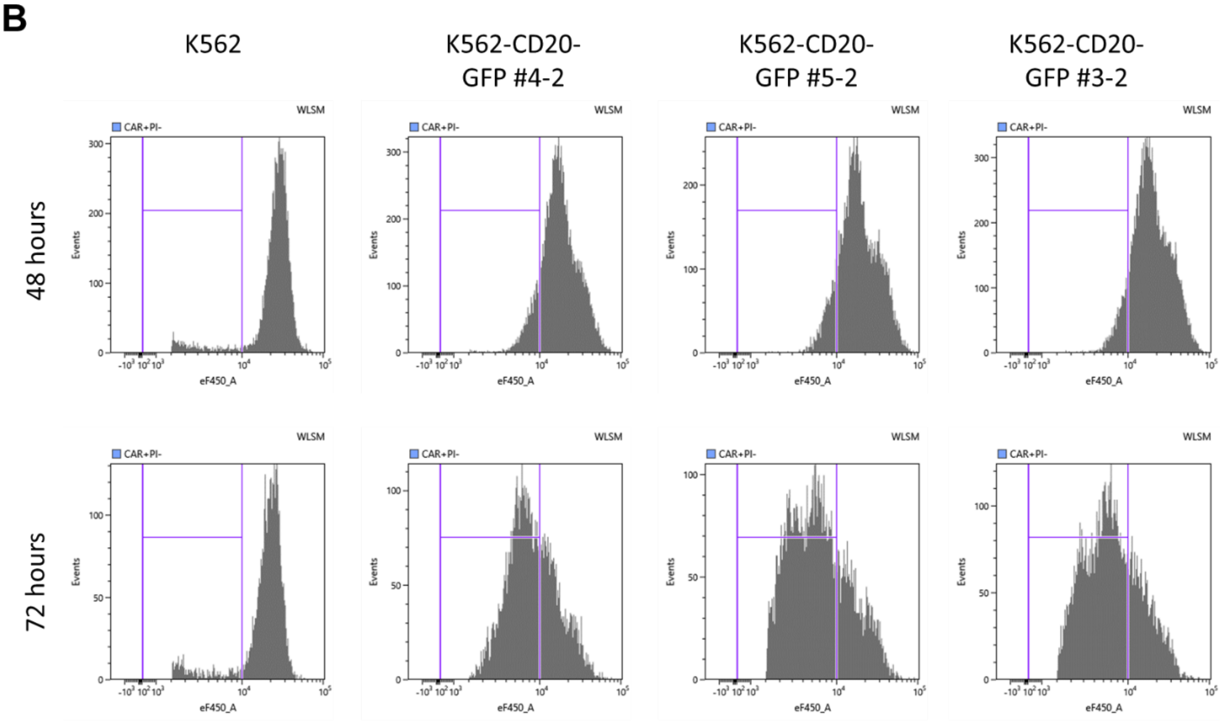

K562
K562-CD20-GFP #4-2
K562-CD20-GFP #5-2
K562-CD20-GFP #3-2
48 hours
72 hours
CAR+PI-
WLSM
Events
eF450_A

**Figure S3**

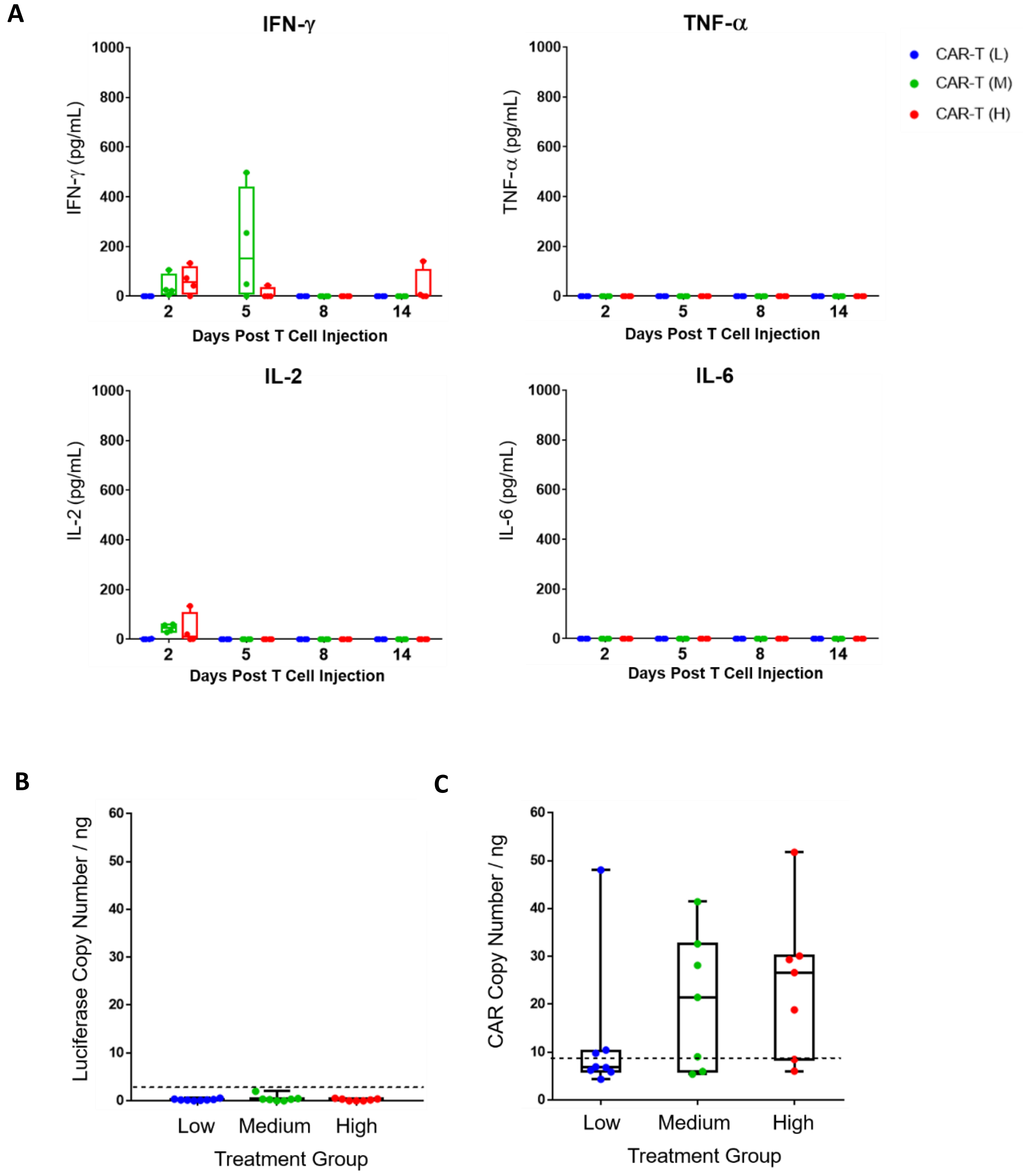
A
IFN-γ
IFN-γ (pg/mL)
TNF-α
TNF-α (pg/mL)
IL-2
IL-2 (pg/mL)
IL-6
IL-6 (pg/mL)
Days Post T Cell Injection
CAR-T (L)
CAR-T (M)
CAR-T (H)
B
Luciferase Copy Number / ng
C
CAR Copy Number / ng
Low
Medium
High
Treatment Group

**Figure S4**

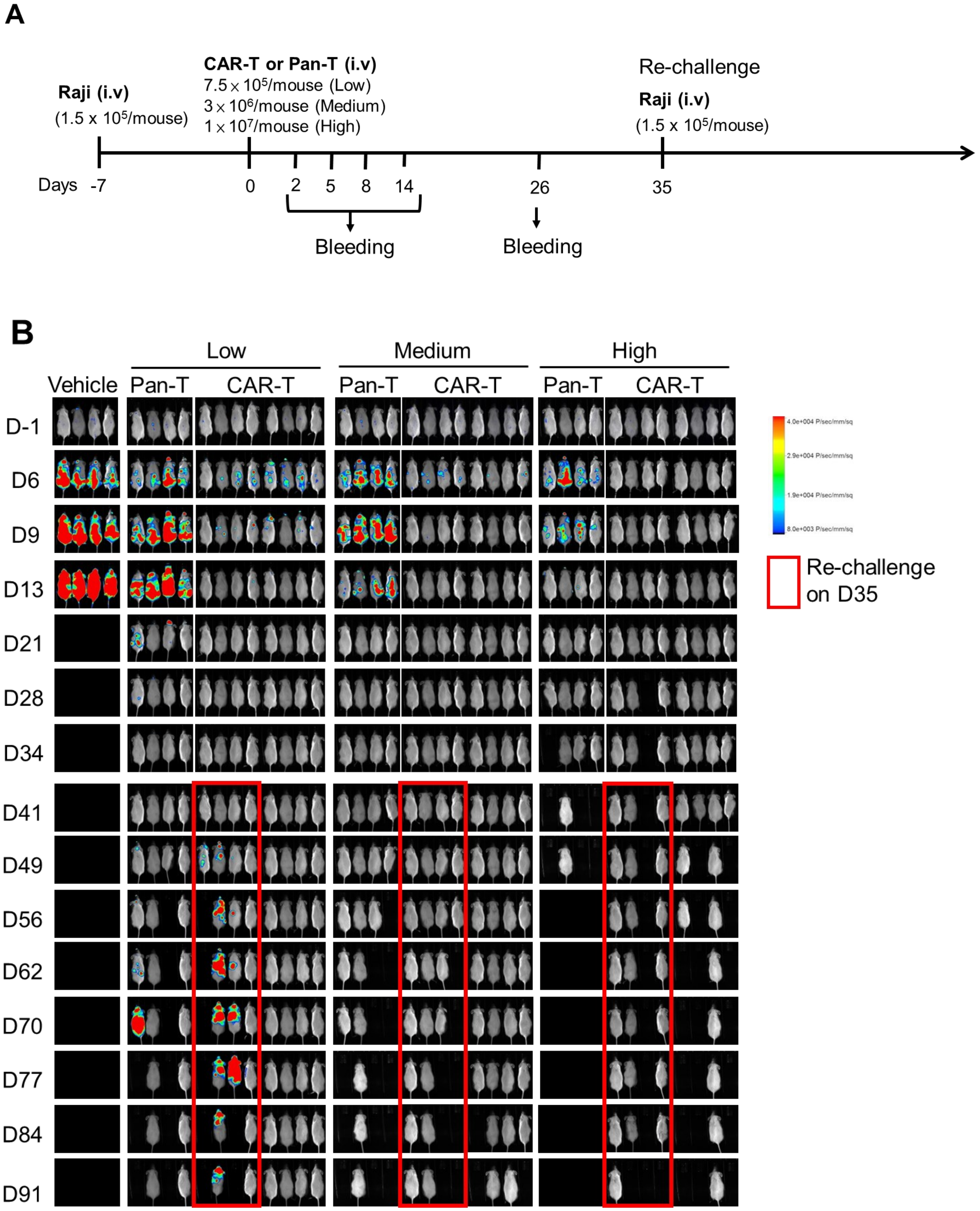
A
Raji (i.v)
(1.5 x 10⁵/mouse)
CAR-T or Pan-T (i.v)
7.5 x 10⁵/mouse (Low)
3 x 10⁶/mouse (Medium)
1 x 10⁷/mouse (High)
Re-challenge
Raji (i.v)
(1.5 x 10⁵/mouse)
Days -7
0
2
5
8
14
26
35
Bleeding
Bleeding
B
Low
Medium
High
Vehicle
Pan-T
CAR-T
Pan-T
CAR-T
Pan-T
CAR-T
D-1
D6
D9
D13
D21
D28
D34
D41
D49
D56
D62
D70
D77
D84
D91
Re-challenge on D35

**Figure S5**

**A**

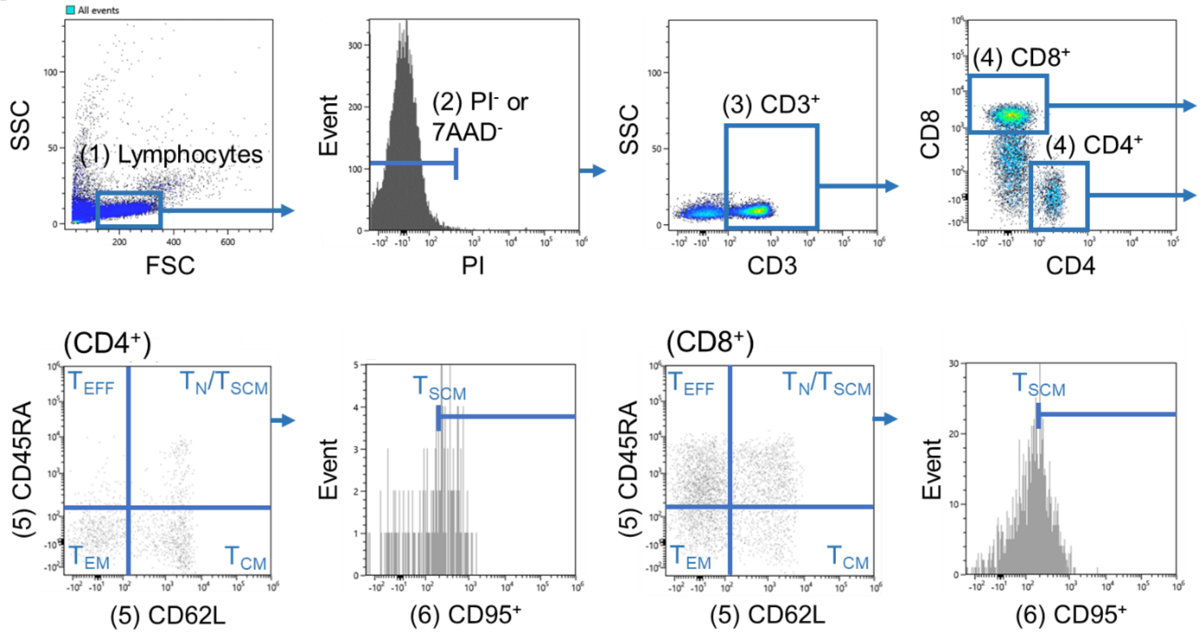
All events
SSC
(1) Lymphocytes
FSC
Event
(2) PI- or 7AAD-
PI
SSC
(3) CD3+
CD3
CD8
(4) CD8+
(4) CD4+
CD4
(CD4+)
(5) CD45RA
TEFF
TN/TSCM
TEM
TCM
(5) CD62L
Event
TSCM
(6) CD95+
(CD8+)
(5) CD45RA
TEFF
TN/TSCM
TEM
TCM
(5) CD62L
Event
TSCM
(6) CD95+

**B**

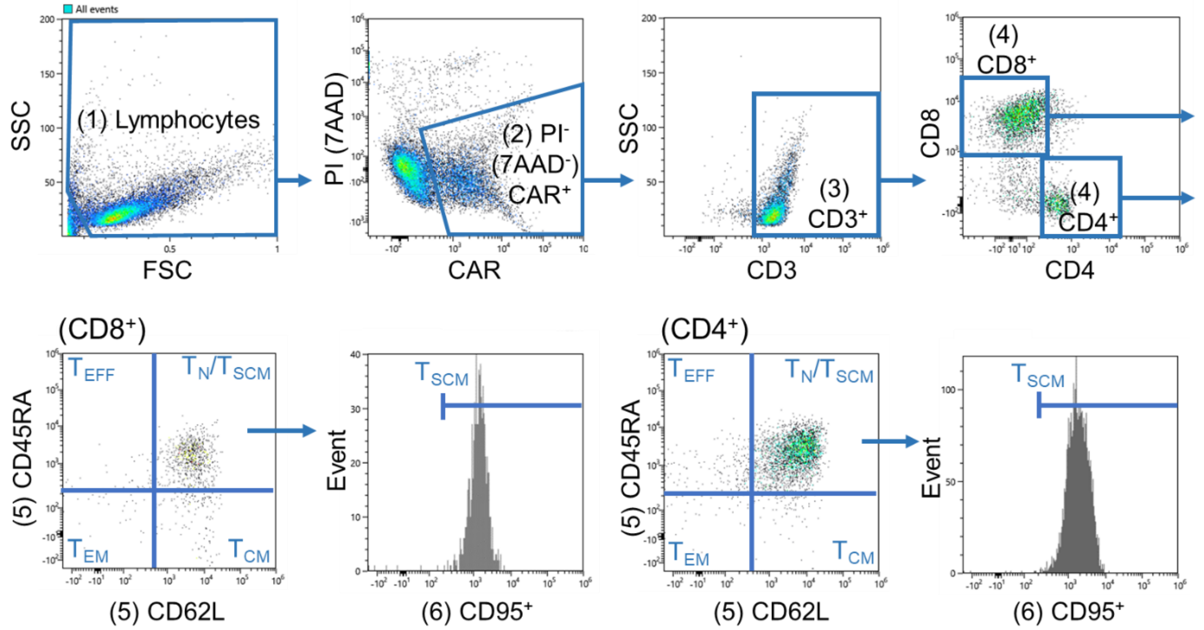
All events
SSC
(1) Lymphocytes
FSC
PI (7AAD)
(2) PI- (7AAD-) CAR+
CAR
SSC
(3) CD3+
CD3
CD8
(4) CD8+
(4) CD4+
CD4
(CD8+)
(5) CD45RA
TEFF
TN/TSCM
TEM
TCM
(5) CD62L
Event
TSCM
(6) CD95+
(CD4+)
(5) CD45RA
TEFF
TN/TSCM
TEM
TCM
(5) CD62L
Event
TSCM
(6) CD95+

**Figure S6**

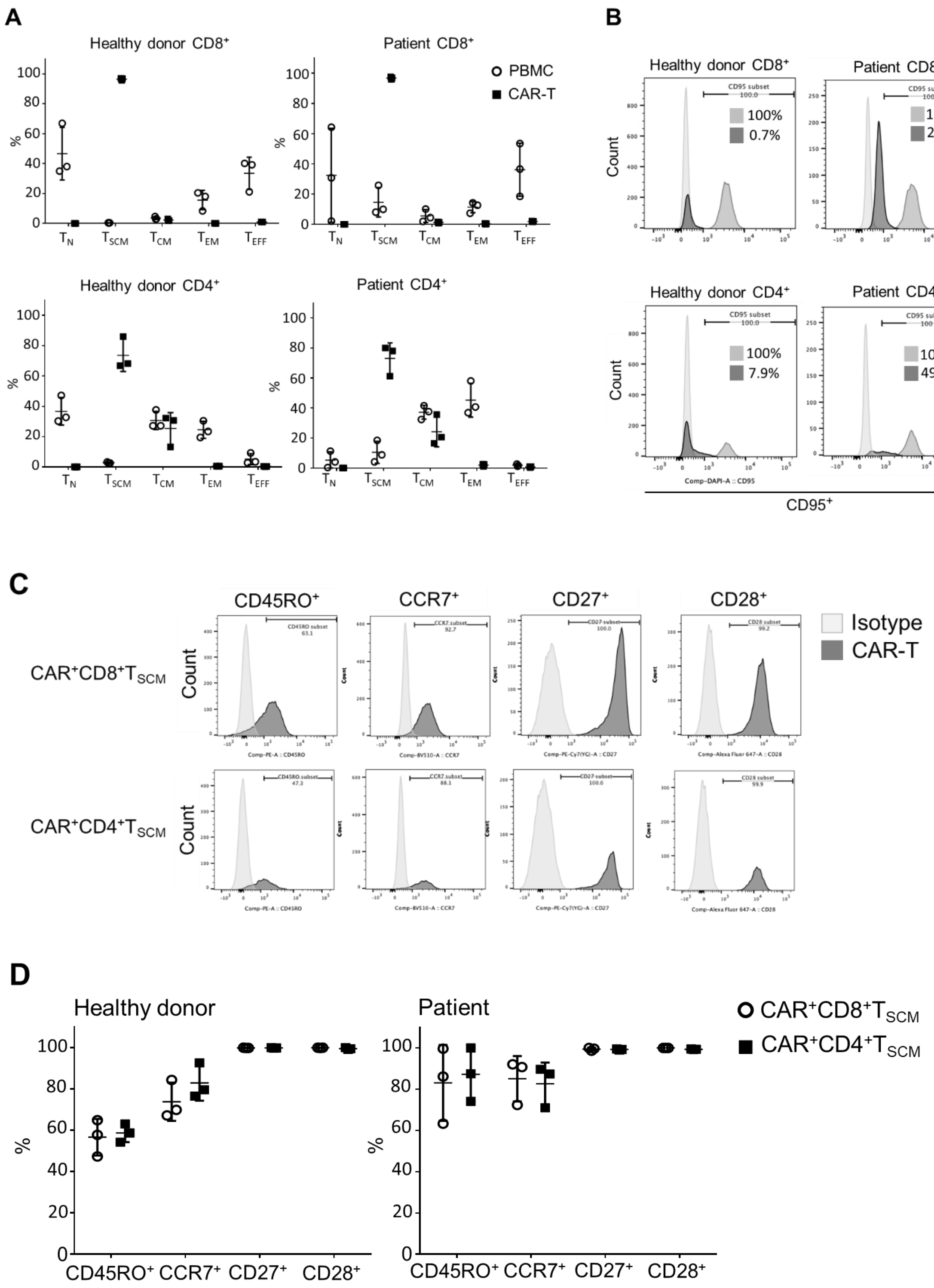
A
Healthy donor CD8⁺
Patient CD8⁺
PBMC
CAR-T
Healthy donor CD4⁺
Patient CD4⁺
B
Healthy donor CD8⁺
Patient CD8⁺
Isotype
CAR-T
PBMC
100%
0.7%
100%
24.3%
Healthy donor CD4⁺
Patient CD4⁺
100%
7.9%
100%
49.6%
Count
CD95⁺
C
CD45RO⁺
CCR7⁺
CD27⁺
CD28⁺
Isotype
CAR-T
CAR⁺CD8⁺TSCM
CAR⁺CD4⁺TSCM
Count
D
Healthy donor
Patient
CAR⁺CD8⁺TSCM
CAR⁺CD4⁺TSCM
CD45RO⁺ CCR7⁺ CD27⁺ CD28⁺

**Figure S7**

**A**

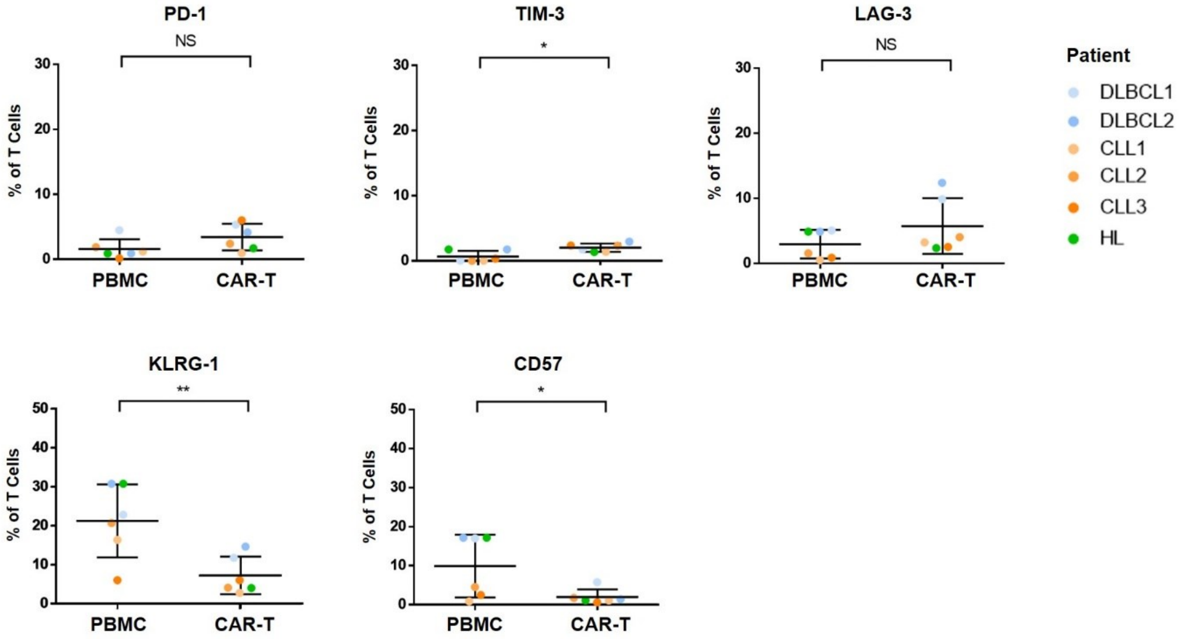
PD-1
NS
% of T Cells
PBMC
CAR-T
TIM-3
*
LAG-3
NS
Patient
DLBCL1
DLBCL2
CLL1
CLL2
CLL3
HL
KLRG-1
**
CD57
*

**B**

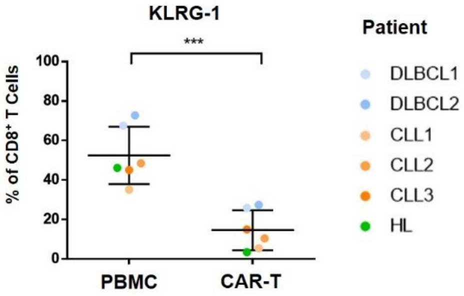
KLRG-1
***
% of CD8⁺ T Cells
PBMC
CAR-T
Patient
DLBCL1
DLBCL2
CLL1
CLL2
CLL3
HL

**Table S1**

| Product | Produced using | CAR-T performance | | | | |
|---|---|---|---|---|---|---|
| | | % of $CAR^+$ | Fold increase | CD8/CD4 Ratio | % $T_N/T_{SCM}$ in $CD4^+CAR^+$ | % $T_N/T_{SCM}$ in $CD4^+CAR^+$ |
| CAR-T* | *qCART*™ | 31.3 | 415.9 | 3.7 | 83.9 | 95.0 |
| | Lentivirus approach | 36.7 | 63.8 | 1.4 | 19.3 | 58.3 |

* Targets to a pan-cancer antigen expressed on NCI-N87 gastric carcinoma cells
